# Mutualisms within light microhabitats drive sensory convergence in a mimetic butterfly community

**DOI:** 10.1101/2024.08.16.607924

**Authors:** J. Benito Wainwright, Theodora Loupasaki, Francisco Ramírez, Iestyn L. Penry Williams, Sam J. England, Annalie Barker, Joana I. Meier, Martin J. How, Nicholas W. Roberts, Jolyon Troscianko, Stephen H. Montgomery

**Affiliations:** School of Biological Sciences, University of Bristol; Bristol, UK; Museo de Zoología QCAZ, Laboratorio de Entomología, Escuela de Ciencias Biológicas, Pontificia Universidad Católca del Ecuador, Quito, Ecuador; Department of Evolutionary Morphology, Museum für Naturkunde-Leibniz Institute for Evolution and Biodiversity Science; Berlin, Germany; Department of Zoology, University of Cambridge; Cambridge, UK; Tree of Life Programme, Wellcome Sanger Institute; Hinxton, UK; Centre for Ecology & Evolution, University of Exeter, Penryn Campus; Penryn, UK

## Abstract

Niche partitioning within variable habitats can expose species to distinct sensory information. Vision is the primary sensory modality used by many animals to interact with their habitat. However, within diurnal terrestrial ecosystems, little is known if, and how, variation in light environments impact species assemblages and visual system evolution. By studying a diverse, sympatric community of mimetic butterflies, we demonstrate that forest architecture creates a mosaic of light microhabitats that drive adaptive sensory convergence and divergence in both peripheral and central sensory systems. Our study provides insights into the dynamic response of visual systems when confronted with similar ecological challenges, and illustrates the wide-reaching consequences of interspecific mutualisms, such as mimicry, on organismal evolution.

## Introduction

Environmental variability is associated with increased biodiversity via adaptive partitioning across multiple ecological dimensions (*1*). This partitioning exposes species to contrasting ecological challenges, including different light environments (*2*), which can impact the efficacy of behaviors such as foraging, mating and predator avoidance (*3*). In aquatic ecosystems, variation in the light environment, such as those caused by spectral or intensity depth gradients, has been inferred to promote ecological diversification with consequences for adaptive visual system evolution (*4–6*). However, the sensory complexity and adaptive influence of light variation in most terrestrial environments, especially in highly diverse tropical rainforests, is less well characterized. As such, the universal role of light variation in shaping the adaptive evolution of community structures is largely unknown.

Evolutionary radiations of Neotropical butterflies provide a powerful system for understanding how the light environment drives both niche partitioning and sensory system evolution in terrestrial ecosystems (*7*). For example, within the 26-million-year-old tribe Ithomiini (∼400 species, Nymphalidae: Danainae), unpalatable species have converged in wing color pattern, morphology, and behavior to increase the efficiency of their warning signal to predators, a mutualistic phenomenon called Müllerian mimicry (*8–12*). Microhabitat segregation of predators allows multiple ‘mimicry rings’ to co-exist within single forest communities as mimetic convergence is associated with convergence in habitat use (*13–15*). Ithomiini also show strong patterns of variation across a range of visual traits, including at the peripheral (compound eye structure), central (brain structure) and molecular (visual pigment coding sequence) level, suggesting lability in sensory system evolution across sympatric species with relatively subtle differences in habitat preference (*16,17*). However, quantitative evaluations of light environments within their forest habitats are lacking, and an absence of dense phylogenetic sampling limits our understanding of the selection pressures shaping the evolution of visual systems in Ithomiini.

Here, we studied a diverse Ecuadorean community of 54 sympatric ithomiine butterfly species, and the lowland rainforest that they live in. We first characterized heterogeneity in sensory microhabitats, and then asked how adaptive species assemblages were strengthened and maintained by abiotic (e.g. changes in the spectral properties of the light environment) and biotic (e.g. mimicry) interactions. Our data provide evidence that ecological interactions among species that occupy a mosaic patchwork of sensory conditions can drive predictable patterns of visual system evolution.

## Materials and Methods

### Field site, animal collection and identification

All fieldwork was conducted along designated trails which surround the Estación Científica Yasuní (0°40’27” S, 76°23’50” W), at Parque Nacional Yasuní, Orellana Province, Ecuador, a ∼4.5 km^2^ area of primary lowland Amazon rainforest where local ithomiine diversity is high (∼60 recorded species) (*13*). All ecological measurements and eye samples were taken in August-October 2022 between 8:00AM (0800h) and 2:00PM (1430h), when butterflies are most active, under permit collection no. MAAE-ARSFC-2021-1763 and export permit no. 023-2022-EXP-IC-FAU-DBI/MAAE. Brain samples were collected in November/December 2011 and September/October 2012 under permit collection no. 0033-FAU-MAE-DPO-PNY and export permit nos. 001-FAU-MAE-DPO-PNY and 006-EXP-CIEN-FAU-DPO-PNY. Permits were obtained from Parque Nacional Yasuní, Ministerio del Ambiente, La Dirección Provincial de Orellana, with support from the Pontificia Universidad Católica del Ecuador (PUCE) and staff at the Estación Científica Yasuní.

Studying a single diverse community allows for robust comparative analysis between species and mimicry rings by eliminating variation due to geographical variables such as altitude and climate. Collecting data across three ∼two-month field seasons also meant that the relative species abundances in our datasets were representative of their natural numbers in the community. Across the three field seasons, 54 ithomiine species were sampled pertaining to the eight local mimicry rings (‘agnosia’, ‘aureliana’, ‘confusa’, ‘eurimedia’, ‘hermias’, ‘lerida’, ‘mamercus’ and ‘mothone’; Fig. 2A; fig. S3) (*13,14*), and nine of the ten major ithomiine subtribes (Dircennina, Godyridina, Ithomiina, Mechanitina, Melinaeina, Methonina, Napeogenina, Oleriina, Tithoreina) (*9*). Genera were identified using wing venation patterns (*31*) and then identified to species level using ID sheets for the races found locally at Yasuní, provided by Dr Keith Willmott, and sexed. The wings of all sampled individuals were kept in transparent envelopes as voucher specimens. Body length (cm) was also measured as a condition-independent measure of body size. In 2022, a maximum of twelve individuals per species were sampled for eye physiological and anatomical analysis. Non-retained individuals were IDed, sexed, and marked on one wing with black permanent marker prior to release to avoid future resampling.

### Spectral and ecological data

#### Transect measurements

To investigate whether changes in forest structure created distinct light environments, spectral measurements were obtained at designated points along the ‘Mirador’ trail, a relatively straight 2.1 km topographically variable transect consisting of nine consecutive ridges (∼250 m above sea level) and valleys (∼240 m above sea level) (Fig. 1). Ten spectral measurements were taken at each ridge/valley replicate at four height categories (1, 2, 3 and 4 m) which were chosen based on flight height data for ithomiines previously reported by Willmott *et al.* (*15*) (Fig. 1). Time of day was also recorded and later binned as ‘early morning’ (0800-0959h), ‘late morning’ (10:00-11:59h) and ‘early afternoon’ (1200-1359h). Individual ithomiine butterflies caught along this transect were assigned to the nearest ridge or valley and one of the four height categories (1, 2, 3 and 4 m), as in Willmott *et al.* (*15*). No butterflies were sampled above 4 m. Although this may have created a sampling bias, other studies have shown ground surveys to provide an accurate proxy of ithomiine vertical stratification and, in practice, very few butterflies were observed flying above 4 m, which was still comfortably within the maximum reach of the hand nets (∼5.5 m) (*8,13,15*).

**Fig. 1.**
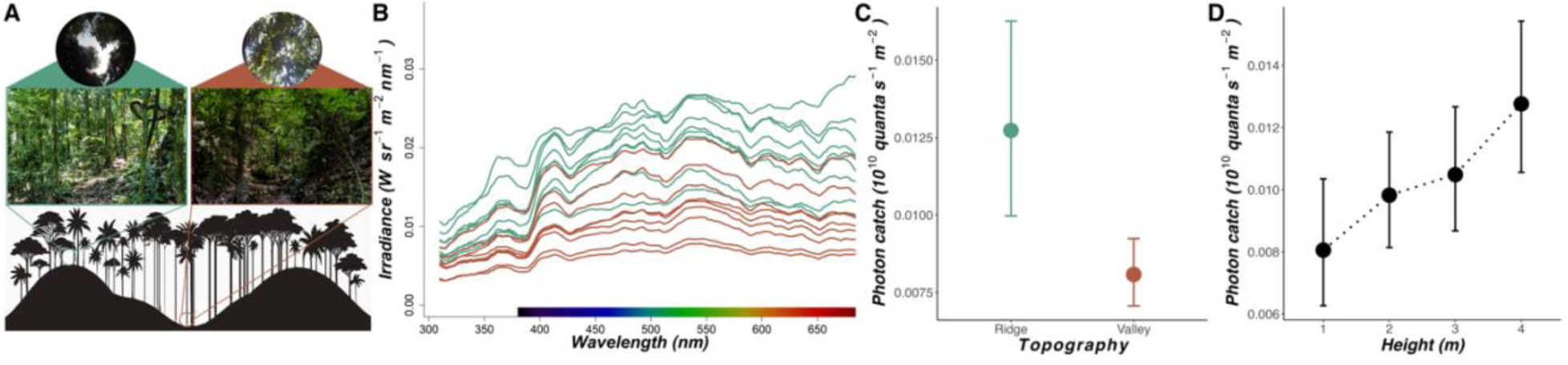
An overview of light environmental differences in tropical rainforests (*N* = 762, 9 ridge/valley replicates). **(A)** Illustration of topographic variation (9 ridges/valleys) along the 2.1 km transect at the field site in Ecuador (bottom), with illustrative digital photographic images from a ridge and valley (middle) and hemispherical, upward facing, 180° fisheye photographs from each location (top). **(B)** Mean raw spectral irradiance values within a 310-700 nm spectral range for each ridge (green) and valley (brown) replicate along the transect. **(C,D)** Mean photon catch (10^10^ quanta s^-1^ m^-2^) for the LW sensitivity function with respect to transect topography (C) and height from the ground (m) (D). Error bars represent 95% confidence intervals.

**Fig 2.**
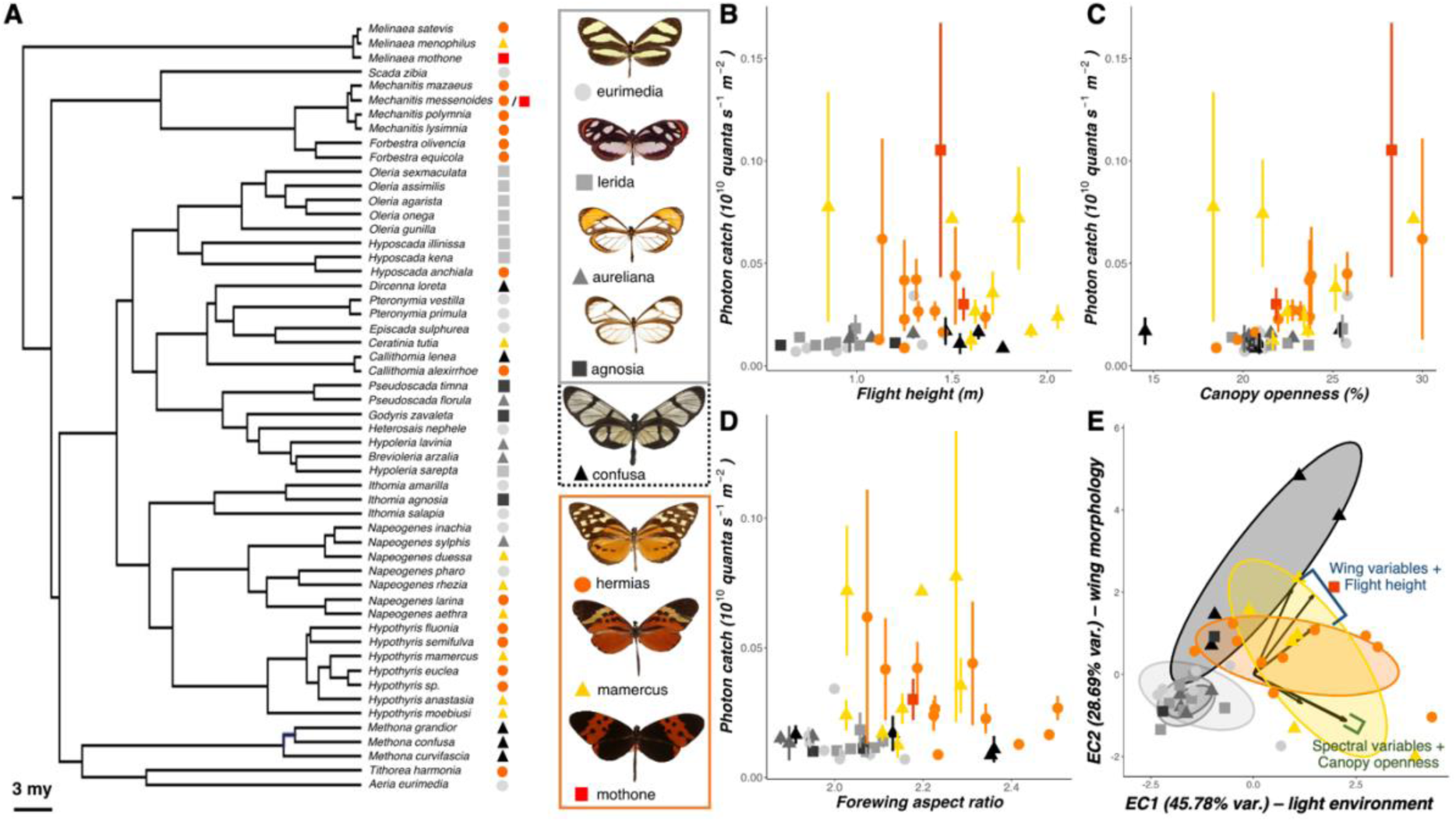
Light microhabitat segregation among ithomiine butterfly mimicry rings. **(A)** A pruned molecular phylogeny [from (*8*)] of the 54 ithomiine species sampled in the Ecuadorean community. Colored symbols at the tips represent the mimicry ring to which each species belongs. Example models of each mimicry ring are shown on the right, grouped based on their general color pattern classification (grey: clearwing, orange: tiger-stripe). ‘Confusa’ has a black dashed border, indicating that it is treated as a separate ‘mimetic cluster’ in later analyses. **(B-D)** Mean photon catch (10^10^ quanta s^-1^ m^-2^) for the LW sensitivity function of each species (*N* = 785, 45 species), coded by mimicry ring, plotted against flight height (m) (B), canopy openness (%) (C) and forewing aspect ratio (D). Error bars indicate standard error. **(E)** Biplot of ecological axes 1 and 2 (EC1 and EC2) from a principal component analysis of mean spectral, ecological, and wing morphological data for each species (*N* = 45 species; see materials and methods). Points and ellipses are coded by mimicry ring. Vector lengths are proportional to the variance at that variable and their loadings are summarized using labelled brackets.

Spectral irradiance measurements were taken using a lightweight, portable, open source, spectroradiometer system (OSpRad), developed by Troscianko (*32*), consisting of a Hamamatsu C12880MA microspectrometer chip (spectral range of 310 to 880 nm, 288-site CMOS sensor, minimal spectral resolution of 9.7 nm), combined with an automated shutter (controlled by a digital servo, Savox SH-0256) and cosine-corrector, and an Arduino Nano microcontroller, contained within custom made, 3D printed ABS plastic housing. OSpRad comes with a custom-built app written in Python and was run via the Pydroid 3 app installed onto a CUBOT Quest Lite smartphone (Android 9.0).

Spectral calibration was performed using a full-spectrum xenon light source (Neewer NW-14EXM) with irradiance measures taken using a Jeti Specbos (1211UV) spectroradiometer with NIST-traceable calibration. OSpRad provided a measure of ambient light intensity from all light sources illuminating the spectroradiometer sensor for each of the 288 sensor photosites (irradiance measured in Wm^-2^nm^-2^; mean integration time: 14.8 ms; no. of scans per measurement: 50; Fig. 1B). For measurements taken at 3 and 4 m, the OSpRad system was attached to the end of hand net poles and raised to the appropriate height.

Hemispherical photography was also used to estimate canopy openness at each ridge/valley replicate by taking an upward facing photograph with the 13 MP camera of a CUBOT Quest Lite smartphone (Android 9.0) attached to a 180° clip-on fisheye lens (Fig. 1A). JPEG images were run through an adapted version of the *hemispheR* R package (*33,34*). The blue channel was extracted from the original RGB images as this provided the best contrast between sky and canopy. Thresholding was then performed using the binarize_fisheye function (using the “Otsu” method), producing binary images analyzed by the gapfrac_fisheye function which calculates the gap fraction for each image within the zenithal angle range of 0-70° using seven zenith angle rings and eight azimuth segments. Canopy openness (%) was then estimated using canopy_fisheye based on the angular distribution of each gap fraction.

#### Individual measurements

Alongside the transect measurements, individual light, and ecological measurements (including hemispherical photography) were taken for 785 butterflies across 45 ithomiine species found along all trails surrounding the Estación Científica Yasuní. Species specific samples sizes varied from 65 for *Hypothyris anastasia* to 1 for *Hypothyris semifulva*, *Ithomia salapia,* and *Napeogenes aethra* with 24 species consisting of more than twelve individuals (Fig. S3). Flight height was estimated for each butterfly using tape measures and the known length of hand net poles. Most butterflies were caught within arm’s reach (∼3 m) but for those that were not, the OSpRad system was attached to the end of hand net poles and raised to the approximate location of initial observation, using distinct features of forest layers and landmarks as reference points.

#### Visual modelling

Irradiance measurements taken by OSpRad were modelled based on the trichromatic visual system of *Danaus plexippus* (Nymphalidae: Danainae), the most closely related species to the Ithomiini for which the maximal sensitivity (λ_max_) of each visual pigment is known (λ_max,,_ UV = 340 nm, B = 435 nm, LW = 545 nm) (*35*). Opsin sequence analysis by Wainwright *et al.* (*17*) suggests that ithomines possess the same number of functional photoreceptors as *D. plexippus*, making *Danaus* a suitable model visual system for calibrating the spectral measurements. The λ_max_ values of these three visual pigments were used to estimate absorbance spectra with nonlinear least-square fitting according to the template by Govardovskii *et al.* (*36*) (fig. S1). Although this template was originally designed to model the spectral sensitivity of vertebrate visual pigments, previous studies have shown that it is practically identical to invertebrate models at accurately describing the normalized sensitivity of rhodopsins, particularly within the visible range (*37*). The absorbance values from these models, and the integration time of each spectral reading, were used to calibrate the raw irradiance measurements. This calculated photoreceptor photon capture (quantum catch, *Q*) for the ultraviolet (UV), blue (B) and long-wavelength (LW) sensitive spectral channels, providing a measure of how many photons are absorbed by each photoreceptor under the given lighting conditions (measured in 10^10^ quanta s^-1^ m^-2^).

It is worth noting that *Danaus,* and likely Ithomiini, also contain a fourth long-wavelength sensitive spectral channel created by the presence of red screening pigment, as indicated by red-reflecting ommatidia in their eyeshine (*17,23*). However, the relative contribution of these screening pigments in shifting the spectral sensitivity of photoreceptors is difficult to disentangle, and the inter-ommatidial connections underlying color processing is poorly understood, making visual modelling using current methods difficult (*22,37*). Our estimated quantum catches are therefore a conservative estimate of the visual information available to an ithomiine butterfly within its respective microhabitat.

A measure of overall photon catch was calculated as the estimated quantum catch of the long-wavelength-sensitive photoreceptor, the primary achromatic channel in invertebrates (*18*). As an alternative, the mean quantum catch of all three spectral channels was also calculated. Both measures of photon catch were log_10_ transformed prior to analysis and analyzed in the same way. To quantify spectral composition, Michelson contrasts were calculated between each spectral channel (*a*) and the summed average of the remaining two spectral channels (*b, c*) ((*Q_a_* – *Q_(b+c)/2_*) / (*Q_a_* + *Q_(b+c)/2_*)), providing an empirical measure of relative ultraviolet, blue, and long-wavelength catch.

Two principal component analyses (PCA) were also conducted for the transect and individual light measurements respectively. In both PCAs, PC1 explained, 98-99% of variation in quantum catch, receiving equal loadings from all three spectral channels, indicating that this is achromatic and equivalent to our measure of overall photon catch (table S1,2). The remaining variance summarized by PC2 and PC3 describe chromatic variation with the UV channel loading negatively against B and LW for PC2, and the B channel loading negatively against UV and LW for PC3. This suggests opponency between these channels, which complements the mechanisms by which colors are processed in other insects (e.g. *39,40)*. In downstream analyses, the distribution of PC values were normalized by log_10_(n + 2) and log_10_(n + 1) transformation for the transect and individual measurements respectively, to avoid log-transformation of negative values. Analyses of these PC axes generally mirrored results from overall photon catch and relative wavelength catch analysis (for full details see table S1,S2).

#### Statistical methods

To test whether the visual environment varied along the transect according to topography, canopy openness, and height from the ground, linear mixed models were constructed using the function lmer in the *lme4* package in R (*34,41*). Both measures of overall photon catch, PC1-3, and relative UV/B/LW catch were each regressed against topography, height (as an ordinal factor), canopy openness, and their interaction. Replicate number, day, and time of day were included as random effects. The significance of each fixed effect was determined by comparing models with and without the variable in question using the *anova()* function.

Based on the results of the above models, we then sought to test whether the abundance of encountered mimicry rings differed between forest microhabitats which displayed significant variation in light environment. This was achieved by constructing Bayesian phylogenetic generalized linear mixed models with the R package *MCMCglmm* (function MCMCglmm, family = “categorical” for binary response variables, family = “gaussian” for continuous, parametric response variables) (*42*) using the inverse correlation matrix of a calibrated and pruned ithomiine phylogeny from Chazot *et al.* (*9*) (packages *phytools* and *ape*) (*43,44*). Default priors were used as fixed effects and uninformative, parameter expanded priors as random effects (G: V = 1,n nu = 1, alpha.mu = 0, alpha.V = 1,000; R: V = 1, nu = 0.002), as in Wainwright and Montgomery (*16*). Species and sex were always included as random effects and models were run for 5,100,000 iterations with a burnin of 100,000. We report the difference in deviance information criterion (ΔDIC) with and without mimicry ring, where lower DICs indicate a better fit and models with ΔDIC < 2 are considered equivalent.

We also performed tests for mimetic segregation between light environments using spectral and ecological measurements obtained from individual butterflies. By applying the same parameters as described above, MCMCglmms were built for each spectral variable with mimicry ring, flight height, canopy openness, and the interaction between these variables included as fixed effects. To assess the significance of continuous variables (flight height and canopy openness), we report the posterior mean (*P*-mean), the 95% CIs, and the *P_MCMC_*.

By calculating species means for each variable of interest, we also recreated all of the above MCMCglmm models using a phylogenetic generalized least-squares regression (PGLS) by implementing the *pgls* function in the *caper* R package (*45*) with Pagel’s λ set to 1 to allow conservative comparisons between models using *anova()*. In this dataset, the polymorphic species *Mechanitis messenoides* was assigned to the “hermias” mimicry ring as this was the observed mimicry pattern for the majority of wild-caught individuals (70.59% ‘hermias’, 29.41% ‘mothone’).

### Summarizing ecological variation

To disentangle the relative roles of light environment and flight-related morphologies in driving visual system evolution (see main text), forewings from individuals sampled in 2011/2012 were photographed dorsally using a DSLR camera and 100 mm macro lens under standardized lighting conditions. Damaged forewings were excluded from the analyses. By running adapted custom scripts from Montejo-Kovacevich *et al.* (*46*) in FIJI (*47*), wing surface area (mm^2^) and aspect ratio were calculated from these images. The latter was defined as the ratio of the major and minor wing axes lengths and was used as a proxy of wing shape, which is known to predict flight speed in many butterfly species (*22*). Wing loading (g mm^-2^) is known to positively predict the lift required to fly at the desired speed and was thus also calculated, by dividing body mass (in this case, the total mass of the head, thorax, and abdomen) by forewing area.

The species means of all wing morphological variables (forewing surface area, aspect ratio, forewing loading) and ecological variables (flight height, canopy openness) were included in a PCA along with the mean quantum catch of the UV, B, and LW spectral channels. The loadings from this PCA revealed all variation to be summarized along two axes; the “light environmental” axis (EC1, 45.99% explained variation) and the “wing morphological” axis (EC2, 28.71% explained variation) (eigenvalue cutoff = 1) which were used in subsequent analyses (see main text; table S3). Mean canopy openness and flight height loaded positively on EC1 and EC2 respectively. Mimetic segregation along both these axes was investigated using PGLS (see ‘Spectral and ecological data – Statistical methods’).

### Physiological and anatomical trait data

#### Eye physiological measurements

Eye physiology was studied at the Estación Científica Yasuní at room temperature using live, wild-caught individuals (*N* = 363, 56.25% female, average of ∼8 individuals per species) with a custom-built ophthalmoscope, previously described by Wainwright *et al.* (*17*), connected to a laptop with the uEye Cockpit program (IDS Software Suite 4.95) installed.

Briefly, butterflies were mounted in slotted plastic tubes, immobilized using plasticine, and oriented with a micromanipulator to face the objective lens, so the frontal region of the compound eye was in view. The ophthalmoscope was then adjusted to focus on the luminous pseudopupil, a region where the optical axes of several ommatidia are aligned and emit colorful eyeshine after dark adaptation, due to the presence of a tapetal reflector at the proximal end of each rhabdom (*48*). The color and heterogeneity of the eyeshine varies between butterfly species but its intensity diminishes quickly upon exposure to light due to intracellular pigment migration towards the rhabdomeres, preventing incoming light from reaching the tapetum (*21,49*). The speed of this pupillary response is known to vary between ithomiine species, suggesting that it might reflect how eyes physiologically respond to spatial and temporal variation in the light conditions present within their respective microhabitats (*17*).

Video recordings of the eyeshine were taken after five minutes of dark adaptation under standard laboratory conditions and imported into FIJI/ImageJ (*43*) where pupillary response time was measured (Fig. 3A). The presence of red-reflecting facets was also noted as this is indicative of screening pigment which is known to create an additional long-wavelength spectral channel in some nymphalid butterflies, the existence of which varies both between and within species (e.g. *21,49)*. We focus on these data as available molecular evidence suggests general conservation of all functioning photoreceptors in ithomiines, which means shifts in eyeshine color are more likely to be a result of changes in screening pigment expression (*17*). The number of reflecting facets were counted and categorized as being yellow or red, from which the ratio of yellow:red reflecting facets was calculated for each individual. The presence of red screening pigments was confirmed in at least one individual of each species.

**Fig. 3.**
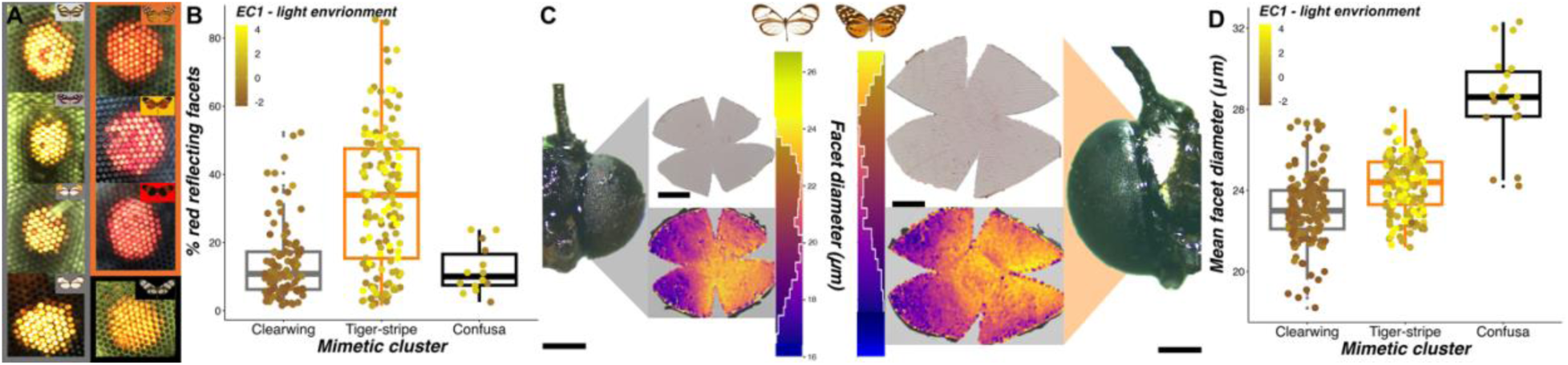
Eye physiological and anatomical associations with light environment and mimicry (*N* = 363, 45 species). **(A)** Example images of the frontal eyeshine from representatives of each mimicry ring within the study community, grouped by mimetic cluster (from left to right: *Ithomia amarilla, Napeogenes larina*, *Oleria gunilla*, *Melinaea menophilus*, *Pseudoscada florula*, *Mechanitis messenoides, Godyris zavaleta*, *Callithomia lenea*). Mimicry ring is indicated in the top right of each panel, with border color denoting mimetic cluster. **(B)** Convergence in the proportion of red reflecting facets (%) for individuals occupying similar light environments, separated by mimetic cluster. **(C)** Example frontal head photographs of *Napeogenes sylphis* (left; mimetic cluster: clearwing) and *Mechanitis mazaeus* (right; mimetic cluster: tiger-stripe) alongside their imaged eye cuticle and its processed output, where individual points denote identified facets, color coded by facet diameter (μm). Scale bars = 500 µm. **(D)** Convergence in mean facet diameter for individuals occupying similar light environments, separated by mimetic cluster. For all boxplots, brown-yellow color shades represent mean EC1 values for each species. Medians (thick horizontal bars), interquartile ranges (boxes), values within 1.5 interquartile ranges of the box edges (whiskers), and possible outliers (datapoints outside whiskers) are plotted.

#### Eye anatomical measurements

Following ophthalmoscopy, heads were removed and preserved in chilled RNAlater (Invitrogen^TM^ AM7021 ThermoFisher Scientific), alongside the remaining body tissue for future molecular work. Samples were kept at 4 °C whenever possible before being returned to the United Kingdom where they were stored at −20 °C. Whole heads were imaged (proboscis and labial palps removed) from the frontal view with LAS X software using a Leica EZ4 W stereo microscope with an integrated 5 MP camera at 20x magnification (Fig. 3C). One compound eye was removed using a fine blade and forceps and placed into 20% sodium hydroxide solution for 3-5 hours to loosen the ommatidia and underlying pigment from behind the eye cuticle. The eye cuticle was then isolated before making four small cuts, two on the dorso-ventral axis and two on the posterior-anterior axis, enough so that it could lay flat. The flattened cuticle was cleaned of debris under 95% ethanol solution and mounted on a clean microscope slide with a small drop of Euparal (Elkay Laboratory Products Ltd) under a cover slip. Mounted slides were then left for a minimum of 24 hours before being imaged with GXCapture software on a Leica M205 C stereo microscope with an integrated 8 MP camera. Images were taken at 1.6x, 2.0x, 2.5x, 3.2x and 4.0x magnification depending on the size of the sample (Fig. 3C).

TIFF images of whole heads were imported into FIJI/ImageJ where inter-ocular distance (mm), defined as the minimum horizontal gap between the two eyes, was measured using the line tool. After correcting the scale in FIJI/ImageJ, facet number, eye surface area, and mean facet diameter were measured from cuticle images using the ommatidia detecting algorithm (ODA) (*51*) (Fig. 3C). This module, written in Python, identifies individual facets from 2D images by extracting periodic signals from each image using a 2D fast Fourier transform. All anatomical variables were log_10_ transformed before any statistical analysis.

#### Sensory neuroanatomical measurements

Relative investment in sensory processing structures was investigated using a separate sample of wild-caught ithomiines (collected in 2011 and 2012) consisting of 392 individuals across 49 species. Species sample sizes varied from 28 for *Hypothyris anastasia* to 1 for *Callithomia alexirrhoe*. *Ceratinia tutia*, *Dircenna loreta*, *Heterosais nephele*, *Ithomia agnosia*, *Pteronymia primula, Pteronymia vestilla*, and *Tithorea harmonia*, with 17 species consisting of more than eight individuals (fig. S3B).

Individuals were dissected at the Estación Científica Yasuní under HEPES-buffered saline (HBS; 150 mM NaCl; 5 mM KCL; 5 mM CaCl2; 25 mM sucrose; 10 mM HEPES; pH 7.4) and fixed in zinc formaldehyde solution (ZnFA; 0.25% [18.4 mM] ZnCl2; 0.788% [135 mM] NaCl; 1.2% [35 mM] sucrose; 1% formaldehyde) for 16-20 hours whilst under agitation. Samples were subsequently washed in HBS three times and placed in 80% methanol/20% DMSO for a minimum of two hours under agitation before being stored in 100% methanol at room temperature and later at −20°C upon arrival to the United Kingdom.

The neuropil volumes within this dataset include those from individuals previously acquired and analyzed by Montgomery and Ott (*52*) and Wainwright and Montgomery (*16*) which were collected during the same two field seasons. The brain tissue of the remaining samples was immunostained against synapsin, a conserved protein expressed at pre-synaptic regions across insects, following the same protocols. Brains were rehydrated in a methanol-Tris buffer series of decreasing concentration (90%, 70%, 50%, 30%, and 0%, pH 7.4), for 10 minutes each, and subsequently incubated in 5% normal goat serum (NGS; New England BioLabs, Hitchin, Hertforshire, UK) diluted in 0.1 M phosphate-buffered saline (PBS: pH 7.4) and 1% DMSO (PBSd) for two hours at room temperature. Samples were stained using anti-SYNORF as a primary antibody (Antibody 3C11; Developmental Studies Hybridoma Bank, University of Iowa, Iowa City, IA; RRID: AB_2315424) in NGS-PBSd, at a dilution of 1:30, and left for 3.5 days under agitation at 4°C. To remove non-bound antibody, three two-hour washes in PBSd were conducted before applying the secondary Cy2-conjugated anti-mouse antibody (Jackson ImmunoResearch; Cat No. 115-225-146, RRID: AB_2307343, West Grove, PA) at a dilution of 1:100 in NGS-PBSd. Samples were incubated for 2.5 days at 4°C under agitation before being subjected to a glycerol dilution series (diluted in 0.1 M Tris buffer, 1% DMSO) of increasing concentration (1%, 2%, and 4% for two hours each, 8%, 15%, 30%, 50%, 60%, 70%, and 80% for one hour each) and then dehydrated in 100% ethanol three times, 30 minutes each. Finally, brains were placed in methyl salicylate and left for ∼30 minutes for the brain tissue to sink and clear, before being replaced with fresh methyl salicylate.

Brains were imaged at the University of Bristol’s Wolfson Bioimaging Facility on a confocal laser-scanning microscope (Leica SP5-AOBS/SP5-II, Leica Microsystem, Mannheim, Germany) fitted with a 10x 0.4 NA objective lens (Leica Material No. 506285, Leica Microsystem). Each sample was scanned from the anterior and posterior side separately using a 488 nm argon laser at 20% intensity, a mechanical z-step of 2 μm, and an x-y resolution of 512×512 pixels. Anterior and posterior image stacks were later merged into a single image stack file in Amira 3D analysis software 2021.2 (ThermoFisher Scientific, FEI Visualization Sciences Group), using a custom *advanced merge* module provided by Rémi Blanc (Application Engineer at FEI Visualization Sciences Group). Prior to image segmentation, the z voxel size of the resulting merged image stack was multiplied by 1.52, to correct for artificial shortening of the z-dimension (*52*). Using Amira 2021.2, every third image of each neuropil of interest was then segmented manually, based on the intensity of the synapsin immunofluorescence, by creating label files for each individual with the *labelfield* module. All intervening unsegmented sections were then assigned to the neuropil of interest by interpolating in the z-dimension, before being edited and smoothed in all three dimensions. Volumetric information was extracted using the *measure statistics* module. This procedure was used to reconstruct the volume (μm^3^) for five of the six primary optic lobe neuropils (medulla, lobula plate, lobula, accessory medulla, and ventral lobula with total optic lobe size being calculated from the sum of these raw volumes). The lamina was not included as it is extremely thin in many smaller species and was easily damaged during the dissections, so could not be obtained reliably for the full range of species. The ventral lobula, a small neuropil found in the optic lobe, was absent in many individuals, particularly females (*17*). Volumes were also reconstructed for the anterior optic tubercle, as the most prominent central brain visual structure, and antennal lobe, as the primary olfactory processing center, to test for comparable effects on olfactory investment. Raw volumes for the anterior optic tubercle and antennal lobe were subtracted from a measure of central brain volume to calculate the “rest of central brain” which acted as an allometric control in subsequent statistical models (Fig. 4A). Each paired neuropil was multiplied by two and all volumes were log_10_ transformed before any analysis.

**Fig. 4.**
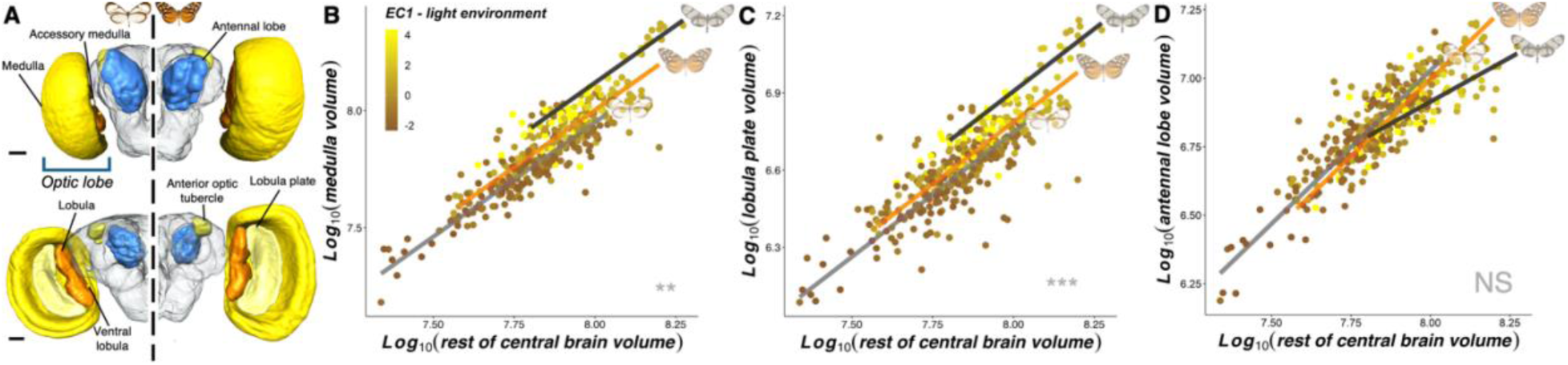
Neuroanatomical associations with light environment and mimicry (*N* = 374 individuals, 40 species). **(A)** Anterior (top) and posterior (bottom) labelled 3D surface reconstructions from the brain of *Ithomia amarilla* (left; mimetic cluster: clearwing) and *Melinaea menophilus* (right; mimetic cluster: tiger-stripe). The figure illustrates all reconstructed neuropils, labelled, and superimposed on an outline of the “rest of central brain”. Scale bars = 100 µm. **(B-D)** Non-allometric convergence between the mean light environment of each species and the level of volumetric investment (μm^3^) in the medulla (B), and lobula plate (C), when scaled against the volume of the “rest of central brain”, with a lack of effect on the antennal lobe (D). All volumetric variables are log10-transformed. Brown-yellow color shades represent mean EC1 values for each species and regression lines for each mimetic cluster, estimated from standardized major axis regressions are superimposed on top. Example models for each mimetic cluster are shown to the right of each line. Asterisks at the bottom right of each panel indicate the significance level of EC1 at predicting relative investment in each neuropil. NS P > 0.05, *P < 0.05, **P < 0.01, ***P < 0.001.

#### Statistical methods – regression analyses

To test whether variation in all our measured visual traits correlate with light microhabitat preference, we regressed each visual trait against EC1 (again, using MCMCglmm, see “Spectral and ecological data – Statistical methods”). EC2, its interaction with EC1, and, where necessary, a relevant allometric control were included in the same model as additional fixed effects. We also tested for covariance amongst physically and functionally connected neuropils in the brain by constructing separate MCMCglmms where neuropils, whose volume showed a significant non-allometric association with either EC axis, were regressed against all other significant neuropils.

In separate models, visual traits were also regressed against individual spectral variables that significantly varied between ithomiine light microhabitats. Light measurements were not taken for individuals sampled for brain tissue, so species means for each spectral variable were used for the analysis of sensory neuropils. We also built separate models where visual traits were regressed against individual wing morphological variables (forewing area, aspect ratio, wing loading) to test whether patterns of visual system evolution are correlated with changes in flight-related wing morphology. Wing morphological variables were collected from individuals sampled for brain tissue only, so species means for each wing variable were used in models involving eye physiological and anatomical traits.

Eye surface area was modelled with inter-ocular distance included as an additional fixed effect to control for allometric effects. To test whether larger eyes have evolved to optimize functional performance by increasing facet size, facet number was modelled with eye surface area as an additional fixed effect. Mean facet diameter was subsequently analyzed without an allometric control to reflect differences in total light capture between species. For models with neuropil volumetric data, “rest of central brain” volume was included as an additional fixed effect to control for allometric effects. These analyses were also recreated using PGLS, as described above, with Pagel’s λ being estimated based on maximum likelihood.

For physiological and anatomical traits which demonstrated evidence of light environment convergence, in both the MCMCglmm and PGLS analysis, we then tested whether mimicry alone could predict convergent shifts in these traits (again, using MCMCglmm and PGLS). For these analyses, mimicry rings were re-clustered based on how species were segregated between light environments. As a post-hoc to the above analysis, we conducted additional tests on anatomical traits which showed convergence with mimicry (in either the MCMCglmm or PGLS analysis), using the sma function in the *smatr* package (*53*). This function detects pairwise shifts in the scaling relationship between the trait of interest and the appropriate allometric control, by testing for slope (β shift), elevation (α or grade shift) and major axis shifts.

#### Statistical methods – evolutionary modelling and analysis of convergence

We used models of morphological evolution to investigate changes in evolutionary dynamics of visual traits that showed evidence of convergence in light environment. We focused on Brownian motion (BM) and Ornstein-Uhlenbeck (OU) models, implemented in the R package *mvMORPH* (*54*). BM models represent random-walk evolutionary processes where the evolutionary rate (parameter σ^2^) is constant. OU processes include both a stochastic BM component and a deterministic tendency towards an optimal value (*θ*), governed by the strength of selection (α) towards adaptive optima, the optimum being the average phenotype toward which lineages subjected to the same ‘selective regime’ have evolved. In the present case, these selective regimes were predefined as the mimetic cluster to which a species belonged. Our hypothesis was that co-mimics, or those sharing similar light environments, will have evolved towards convergent adaptive optima in sensory anatomical trait space.

Prior to analysis, we used the species means dataset to conduct a PCA using visual traits that showed evidence of light environment convergence, plus “rest of central brain” volume as an allometric control where relevant (see main text). The significant axes from this PCA were also regressed against EC1, EC2, and their interaction, using PGLS. The reconstructed history of the selective regime on which each model was fitted, was constructed using 500 character maps in *phytools* using the ‘make.simmap’ function (*44*). These simulated character maps were used to fit BM and OU models with single (BM1, OU1) and regime-specific optima (BMM, OUM) to univariate models (functions ‘mvBM’ and ‘mvOU’). We also constructed an “early burst” (EB) model, where visual systems diversify rapidly early on in cladogenesis (function ‘mvEB’). The fit of the resulting models was compared using the mean corrected Akaike information criterion (AICc) with lower AICc values indicating an improved model fit.

We also quantified levels of evolutionary convergence in sensory morphology between co-mimics, or those sharing similar light environments, using the C indexes (C1-4) calculated within the *convevol* package (*55*). Data were simulated under Brownian motion 1,000 times to gauge significance. Lastly, a phenogram which maps visual trait evolution was created using the “phenogram” function in *phytools* where crossing branches indicate convergent evolution (*44*).

## Results and Discussion

### Variation in ecologically relevant forest light creates a mosaic of microhabitats

We collected spectral irradiance measurements across a 2.1 km topographically variable forest transect, consisting of nine consecutive ridges and valleys, in Parque Nacional Yasuní, Ecuador (Fig. 1A). Photoreceptor spectral sensitivities of the closely related *Danaus plexippus* (Nymphalidae: Daniainae; fig. S1; see materials and methods) were used to estimate the photon catch (defined here as the absorption of light photons captured by the long-wavelength-sensitive photoreceptor (*18*)), and the relative number of photons captured by the ultraviolet (UV), blue (B), and long-wavelength (LW) photoreceptors (see materials and methods). Overall, the architecture of forest ridges created brighter and broader spectrum visual environments than forest valleys (GLMM: photon catch: χ^2^_1_ = 56.890, p < 0.001; Relative B catch, χ^2^_1_ = 17.262, p < 0.001; Relative LW catch, χ^2^_1_ = 11.098, p = 0.001; Fig. 1B,C; fig. S2A; table S1). The amount of light relevant to the butterfly visual system also showed a positive correlation with measurement height (GLMM: χ^2^_1_ = 34.067, p < 0.001; Fig. 1D) and canopy openness (χ^2^_1_ = 11.023, p = 0.001; fig. S2C). No interactions between topography, height from the ground, or canopy openness were found for any other spectral variable (table S1).

To complement our transect data, we recorded spectral irradiance measurements and flight height at the capture location and position of 785 wild, individual ithomiine butterflies (45 species; Fig. 2A; fig. S3). This expanded our sampling of the light environment across a broader area (∼4.5 km^2^) of forest where butterflies were naturally flying. While controlling for potential phylogenetic effects in species’ visual niches, photon catch and the relative abundance of blue wavelengths were both positively correlated with canopy openness. We did not see the same effect for flight height, but this is likely explained by aforementioned topographic effects which decouple flight height from distance to the canopy (MCMCglmm: photon catch, P-mean = 0.010, 95% CI = 0.007-0.013, P_MCMC_ < 0.001; Relative B catch, P-mean = 0.001, 95% CI = 0.001-0.001, P_MCMC_ < 0.001; Fig. 2B,C; table S2A). Hence, the abundance and composition of ecologically important light within tropical forests varies along both vertical (height and topography) and horizontal (canopy openness) axes, validating previous categorizations of “forest shade” and “small gap” light microhabitats (*2*).

### Light microhabitat partitioning among mimicry rings

Although ithomiine mimicry rings differ in topography and height (MCMCglmm; topography (valley vs. ridge), ΔDIC = 7.856; height, ΔDIC = 5.989; fig. S4,5), our spectral irradiance measurements from individual butterflies revealed finer scale microhabitat partitioning between the eight mimicry rings (MCMCglmm: photon catch, ΔDIC = 10.142; Relative LW catch, ΔDIC = 5.053; Fig. 2C,D; table S2A). Pairwise contrasts revealed that all significant comparisons were between mimicry rings with ‘tiger-stripe’ (opaque) and ‘clearwing’ (particularly transparent) color patterns (Fig. 2; fig. S5,S6; see table S2A), suggesting a fundamental ecological split mirroring major color pattern traits. Species involved in brightly colored tiger-stripe mimicry rings flew higher, and occupied brighter, broader spectrum and more variable light environments compared to species involved in clearwing mimicry rings, which were confined to shaded forest at lower elevations. Together, our results demonstrate that heterogeneity in visual niche preference and mutualistic mimetic interactions can co-evolve, generating substantial ecological diversity and maintaining adaptive niche assemblages within tropical forests (*19*). This co-evolution is likely driven, at least in part, by differences in predator communities, which are known to segregate across forest types (*15*), and the adaptive evolution of anti-predator coloration and conspecific signaling (*20*).

### Spectral variation is independent of flight behavior

Ithomiines occupying similar mimicry rings are known to converge in flight-related morphologies, forming three mimetic clusters in flight morphospace within rainforest communities (*9*). In support of this, we found a positive correlation between forewing aspect ratio and light environment (Fig. 2D; fig. S7; table S2B). Therefore, variation in flight speed and performance could conceivably drive changes in visual processing independently of light microhabitat preference (*10,21,22*). To disentangle these effects, we conducted a principal component analysis (PCA) on three major light environmental variables (estimated photon catch for the ultraviolet, blue, and long-wavelength photoreceptors), two ecological variables (flight height and canopy openness), and three flight-related morphological measurements (forewing surface area, aspect ratio, and wing loading; see materials and methods). This resulted in two ‘ecological axes’ (ECs) which explained 45.78% and 28.69% of variation respectively. All spectral variables loaded most positively onto EC1, and all wing morphological variables loaded most positively onto EC2 (Fig. 2E; table S3), providing two independent ‘light environment’ (EC1) and ‘flight related wing morphology’ (EC2) axes which were used in subsequent analyses. ‘Tiger-stripe’ and ‘clearwing’ mimetic clusters segregated along EC1 but the ‘confusa’ mimicry ring also diverged along the EC2 ‘wing morphology’ axis, despite occupying similar light environments to clearwing co-mimics (PGLS: EC1, lambda = 0.000, F_7,37_ = 7.686, p < 0.001; EC2, lambda = 0.000, F_7,37_ = 3.659, p = 0.004) (Fig. 2E, table S3). This confirms previous findings of convergent flight morphologies among clearwing and tiger-stripe ithomiine co-mimics, with the ‘confusa’ mimicry ring forming a third distinct ‘mimetic cluster’ that is treated separately hereafter (*10*).

### Light microhabitat is associated with the evolution of peripheral light reception

We hypothesized that the patterns we observed in light microhabitat preference would impact the selection regimes acting on the visual system. Using wild-caught butterflies, we first examined physiological data extracted from video recordings of eyeshine, which reveal screening pigments that shift the spectral sensitivity of photoreceptors towards longer wavelengths, resulting in red reflecting facets (*17,23*) (Fig. 3A). We found that the relative abundance of red-reflecting facets was positively correlated with EC1, meaning red-reflecting facets were more abundant in species occupying brighter light environments (MCMCglmm: P-mean = −0.114, 95% CI = −0.173 - −0.059, P_MCMC_ = 0.001; Fig. 3B; fig. S8; table S4A). This suggests that shifts in screening pigment expression provide an evolutionary mechanism, operating in conjunction with the spectral tuning of visual pigments, to shape visual specialization (*17*). Eyeshine exposure is followed by a rapid pupillary response, which we used to assay how rapidly eyes physiologically respond to sudden shifts in lighting conditions. No significant correlation was found between EC1 (or EC2) and the speed of this pupillary response (MCMCglmm: P-mean = 0.116, 95% CI = 0.004 – 0.231, P_MCMC_ = 0.054), suggesting that differences in temporal light variability between light microhabitats does not explain variation in this trait.

We subsequently investigated how these eye physiological adaptations have co-evolved with eye structure by measuring the surface area of the eye cuticle, the number of facets, and mean facet diameter (Fig. 3C). Eye surface area showed a strong allometric relationship with inter-ocular distance, here used as an allometric control (fig. S8,9; see materials and methods), but nevertheless showed a significant non-allometric positive association, independently, with respect to both EC1 (light environment) and EC2 (wing morphology) (MCMCglmm: EC1, P-mean = 0.031, 95% CI = 0.019 – 0.043, P_MCMC_ < 0.001; EC2, P-mean = 0.040, 95% CI = 0.024 – 0.055, P_MCMC_ < 0.001; fig. S8,9). Species occupying more illuminated light environments (i.e. species belonging to tiger-stripe mimicry rings) or with forewings equipped for greater flight speeds (i.e. ‘confusa’ co-mimics) have convergently evolved larger eyes. When regressing facet number against eye surface area, we again found independent, non-allometric effects of light environment and wing morphology (MCMCglmm: EC1, P-mean = 0.008, 95% CI = 0.004 – 0.012, P_MCMC_ < 0,001; EC2, P-mean = 0.008, 95% CI = 0.003 – 0.014, P_MCMC_ = 0.004; fig. S8,9; table S4A), suggesting that larger eyes have evolved wider facets for maximizing light capture, as well as larger numbers of facets. This was confirmed by regressions of mean facet diameter against EC1 and EC2 (MCMCglmm: EC1, P-mean = 0.008, 95% CI = 0.004 – 0.011, P_MCMC_ < 0.001; EC2, P-mean = 0.010, 95% CI = 0.005 – 0.015, P_MCMC_ < 0.001; Fig. 3C,D; fig. S8,9; table S4A). Therefore, while it remains to be confirmed how these anatomical shifts impact functional performance, the physical structure of the eye has repeatedly evolved in response to increased light abundance found within more open forest microhabitats.

### Light microhabitat is associated with the evolution of investment in visual brain centers

Next, we followed the trajectory of visual information to assess whether the light environment specifically impacts the peripheral visual system, or also shapes investment in sensory regions of the central brain (Fig. 4A). After accounting for overall brain size (which correlates strongly and positively with other allometric controls; fig. S10), significant non-allometric associations were found between EC1 and the size of optic lobe neuropils, synapse-dense regions which form the primary visual processing center in the brain (*24*). In these analyses, investment in the optic lobe as a whole (MCMCglmm: P-mean = 0.023, 95% CI = 0.013-0.033, P_MCMC_ < 0.001), and three of the largest visual neuropils (the medulla, lobula plate and lobula; Fig. 4B,C; fig. S11; table S4A,S4B), were largest in species found in more illuminated microhabitats. An independent, positive, non-allometric effect of EC2 was also found for the scaling of the same neuropils (fig. S12; table S4A,S4B). After controlling for covariance between these three physically and functionally connected neuropils, significant effects of EC1 and EC2 were retained for the lobula plate (MCMCglmm: EC1, P-mean = 0.010, 95% CI = 0.004 – 0.016, P_MCMC_ = 0.001; EC2, P-mean = 0.011, 95% CI = 0.003 – 0.019, P_MCMC_ = 0.007; table S4A), suggesting that selection shaping behavioral processes primarily mediated by this structure might particularly explain volumetric shifts in the optic lobe.

Similar associations were not found for sensory neuropils within the central brain, including the antennal lobe, the primary olfactory processing center, implying that greater investment in visual processing does not correlate with reductions in olfactory investment, or *vice versa* (*25*) (Fig. 4D; fig. S11; table S4A,B). Previous data showing extensive visual system variation from insectary-reared individuals of four ithomiine species also suggests that these shifts are likely heritable, rather than the result of environmentally-induced plasticity (*16,26*). Our results therefore demonstrate that both peripheral and central components of the visual pathway have evolved to exploit favorable abundance of visual information present in their specific environment. This contrasts with some previously reported patterns of evolution in Lepidoptera where visual systems evolve to maximize the capture of unfavorable light abundance, as is seen during transitions to nocturnality in many species (e.g. *27)*.

### Variation in light environments drives sensory convergence among co-mimics

In Ithomiini, and other radiations of mimetic butterflies, shifts in mimicry patterns are known to instigate speciation processes, with ecological segregation accelerating reproductive isolation (*28–30*). As such, secondary local adaptation across distinct environmental conditions may be a critical factor in the diversification of species. Given our evidence that multiple aspects of variation in the visual system are associated with light environments, we next tested whether ecological convergence among co-mimics could also predict adaptive convergence in sensory traits. We approached this through multiple statistical methods. First, when data were analyzed at an individual level using phylogenetic generalized linear mixed models, the addition of mimetic cluster (three groups: tiger-stripe, clearwing, confusa) improved model fit in all measured visual traits, except for the relative abundance of red-reflecting facets (Fig. 3,4; table S5).

Standardized major axis regressions also confirmed that these differences were the result of non-allometric adaptive ‘grade shifts’ (shifts along the y-axis) between mimetic clusters, except for the lobula which showed a significant shift in the allometric slope (Fig. 3,4; table S5). These results were further supported by a significant effect of mimetic cluster on all visual traits when species means were analyzed with phylogenetic generalized least squares regression (PGLS), with the exception of eye surface area and facet number (see table S5).

These findings demonstrate that mutualistic mimetic interactions lead to sensory convergence due to indirect effects of niche similarity. To test whether this convergence arose via adaptive evolutionary processes, we summarized variation in traits that showed significant non-allometric light environmental effects (relative abundance of red-reflecting facets, eye surface area, facet number, facet diameter, and the volume of the medulla, lobula plate, and lobula, and “rest of central brain” volume as an allometric control) along a single principal component axis (which explained 78.95% of variation in the data). Although this axis might capture allometric and non-allometric components of trait variation, the loading of our allometric control (“rest of central brain” volume) along this axis is relatively low compared to other traits (table S6) suggesting non-allometric differences are a more distinguishing feature of visual structural variation. With this summary component, which reflects variation across the visual pathway and correlates independently with both EC1 and EC2 (PGLS, lambda = 1.000; EC1, t = −5.864, p < 0.001; EC2, t = −7.768, p < 0.001), we constructed single and multi-peak Brownian motion (BM), Ornstein-Uhlenbeck (OU) and early burst (EB) models of evolution (see materials and methods). EC1 (light environment) and EC2 (flight-related wing morphology) were also modelled in the same way. Interspecific variation in light environment preference, flight-related wing morphology, and variation in the visual pathway, consistently fitted multi-peak OU models above all other models, indicating that species in the same mimicry ring are attracted towards convergent adaptive optima (Fig. 5; table S7). Visual system convergence was further supported by equivalent models constructed for eye and brain structures separately (using separate eye and brain PC axes) and by additional tests for evolutionary determinism (fig. S13; table S7). Together, these analyses provide strong evidence that directional evolutionary change, driven by convergence in both light microhabitat and flight-related wing morphology, has generated similarities in a suite of visual system traits, reflecting local adaptation that may support the maintenance of niche partitioning and, potentially, create reproductive barriers within the Ithomiini radiation.

**Fig. 5.**
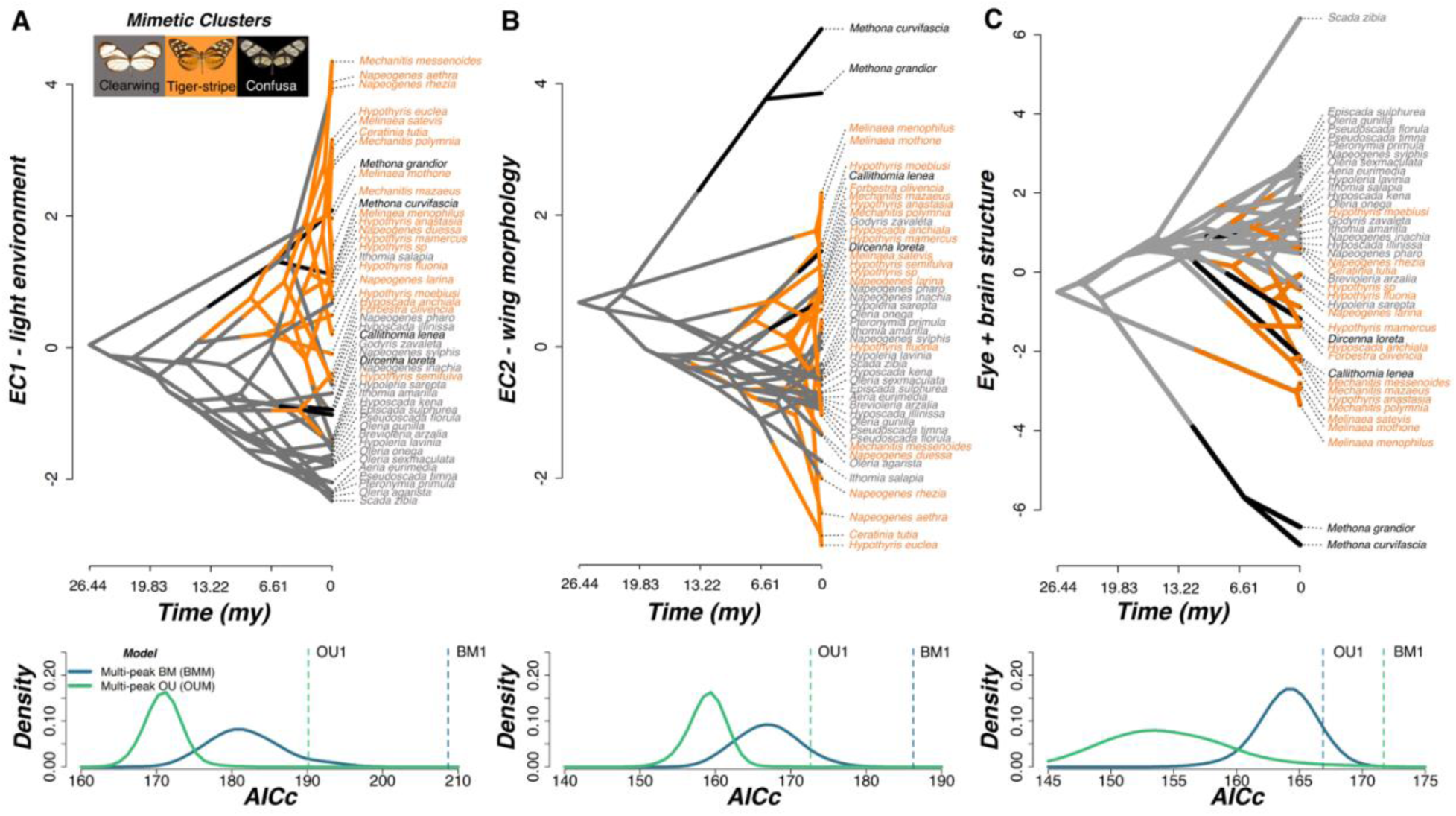
Adaptive convergence in light environment, wing morphology, and visual system structure among co-mimics. **(A-C)** Phenograms (top panel) based on EC1, EC2 (A,B) (*N = 45* species), and PC1 of a principal component analysis which summarized variation in eye and brain structure based on traits which showed significant light environmental effects (C) (*N = 40* species). Instances where branches cross and concentrate in a given area indicate convergent lineages. Below each phenogram are Kernel density plots of small sample corrected Akaike Information Criterion (AICc) scores obtained from multi-peak Brownian motion (BMM) and Ornstein-Uhlenbeck (OUM) evolutionary models, constructed from 500 simulated character maps where species belonging to the same mimetic cluster were assigned to the same selective regime. Blue and green vertical dashed lines indicate the AICc score for single-peak BM (BM1) and OU (OU1) models respectively.

### Summary

Our study provides empirical evidence that relatively small shifts in terrestrial light environment properties can shape adaptive and predictable visual system evolution between sympatric species, over very small spatial scales. Our data also offers insights into how variation in ecologically relevant light environments shapes the assemblage of diverse communities. Specifically, we have shown that light abundance and spectral composition within rainforests can contribute to multiple dimensions of niche separation that overcome phylogenetic niche conservatism. We have further demonstrated how these fluctuations in ambient light conditions can be a potent selective force, predictably driving visual systems towards distinct adaptive peaks. Notably, our data also reveal how mutualistic mimetic interactions can have broad-reaching and perhaps surprising effects on organismal evolution, by indirectly impacting multiple levels of visual pathways. We conclude that ambient light plays a crucial role in shaping both tropical forest communities and patterns of visual system evolution. Together, this introduces mimetic butterflies as an informative model system for disentangling the many manifestations by which species have evolved to perceive and process their sensory world.

## Acknowledgments

We are extremely grateful to J. C. Armijos, Á Barragán, P. Jarrín, D. Lasso, E. Moreno, and C. Padilla from the Estación Científica Yasuní, Pontificia Universidad Católica del Ecuador (PUCE), M. Arévelo and M. F. Checa for assisting with the acquisition of collection and export permits, and K. Willmott for his identification sheets and assistance with species identification in 2022. We thank G. Montejo-Kovacevich for support on forewing measurements, S. Wright for providing guidance on the eye dissection protocol and W. Hurley for assisting with imaging mounted cuticle samples. We also thank K. Jepson, M. Jepson, and colleagues at the University of Bristol’s Wolfson Bioimaging Facility for confocal microscope support, E. Martin-Silverstone from the University of Bristol’s XTM Facility for Amira support, and N. Chazot for providing permission to use his phylogenetic tree. We acknowledge the use of ChatGPT Pro 3.5 (Open AI, https://chat.openai.com) for assisting in the creation of Fig. 1A and thank K. Manser for assistance. Lastly, we thank R. M. Merrill and L. H. Yusuf for insightful comments on a draft manuscript.

## Funding

Natural Environment Research Council Independent Research Fellowship NE/N014936/1(SHM)

Natural Environment Research Council GW4+ Doctoral Training Partnership studentship (JBW)

1851 Royal Commission for the Exhibition Research Fellowship (SHM) Royal Society Small Grant RG110466 (SHM)

Leverhulme Trust Early Career Research Fellowship (SHM)

Natural Environment Research Council Independent Research Fellowship NE/P018084/1 (JT)

Air Force Research Laboratory/Air Force Office of Scientific Research through the European Office of Aerospace Research and Development grant FA9550-18-1-7005 (NWR)

1851 Royal Commission for the Exhibition Research Fellowship (JBW)

## Author contributions

Conceptualization: JBW, SHM

Methodology: JBW, ILPW, SJE, MJH, NWR, JT, SHM

Investigation: JBW, TL, FR, ILPW, SJE, AB, JIM, SHM

Visualization: JBW, SJE, SHM

Funding acquisition: JBW, SHM

Project administration: JBW, SHM

Supervision: NWR, JT, SHM

Writing – original draft: JBW, SHM

Writing – review & editing: All authors

## Data and materials availability

All raw data, R analysis code, uncalibrated spectral data, visual modelling templates, hemispherical photography images, forewing images, eyeshine video recordings, eye cuticle images and analyses output, confocal microscopy image stacks, and 3D brain reconstruction files are available from Zenodo: https://tinyurl.com/348wst9w

## Supplementary Materials

*Figs. S1 to S13*

- **Fig. S1:** The butterfly visual model used to calibrate raw spectral irradiance values (p. 13)
- **Fig. S2:** Additional ecological associations with light environment across a topographically variable forest transect (p. 14)
- **Fig. S3:** Ithomiini species abundances within the Ecuadorean community (p. 15)
- **Fig. S4:** Relative mimicry ring abundance at ridge and valley sites along a topographically variable forest transect (p. 16)
- **Fig. S5:** Flight height (m) differences between ithomiine species, coded by mimicry ring (p. 17)
- **Fig. S6:** Variation in spectral composition between ithomiine microhabitats (p. 18)
- **Fig. S7:** Forewing aspect ratio differences between ithomiine species, coded by mimicry ring (p. 19)
- **Fig. S8:** Additional eye physiological and anatomical associations with light environment and mimicry (p. 20)
- **Fig. S9:** Eye anatomical associations with wing morphology (p. 21)
- **Fig. S10:** Positive correlations between log10-transformed allometric controls (p. 22)
- **Fig. S11:** Additional neuroanatomical associations with light environment and mimicry (p. 23)
- **Fig. S12:** Neuroanatomical associations with wing morphology (p. 24)
- **Fig. S13:** Convergence in eye and brain structure separately among co-mimics (p. 25)

**Other Supplementary Materials for this manuscript include the following:**

***Wainwright et al_Supplementary_tables.xlsx: Tables S1 to S7 (details on p. 26)***

- **Table S1:** Light microhabitat variation and partitioning along a topographically variable transect.
- **Table S2A**: Light microhabitat variation and partitioning for individually caught ithomiine butterflies – MCMCglmm and PGLS segregation analyses.
- **Table S2B:** Light microhabitat variation and partitioning for individually caught ithomiine butterflies – spectral PCA and light-wing morphological associations.
- **Table S3:** Ecological principal component analysis (PCA).
- **Table S4A:** Visual system associations with light environment and flight-related morphology – ecological PC axes.
- **Table S4B:** Visual system associations with light environment and flight-related morphology – spectral and wing morphological variables.
- **Table S5:** Sensory convergence between co-mimics.
- **Table S6:** Visual system principal component analyses (PCA).
- **Table S7:** Evolutionary modelling of convergence.

***Wainwright et al_Supplementary_data.xlsx: Data S1 to S5 (details on p. 28)***

- **Data S1:** Transect spectral and ecological data
- **Data S2:** Individual spectral and ecological data
- **Data S3:** Individual eye physiological and anatomical data
- **Data S4:** Individual neuroanatomical data
- **Data S5:** Summary data: Species means of all variables

---

**Fig. S1.**
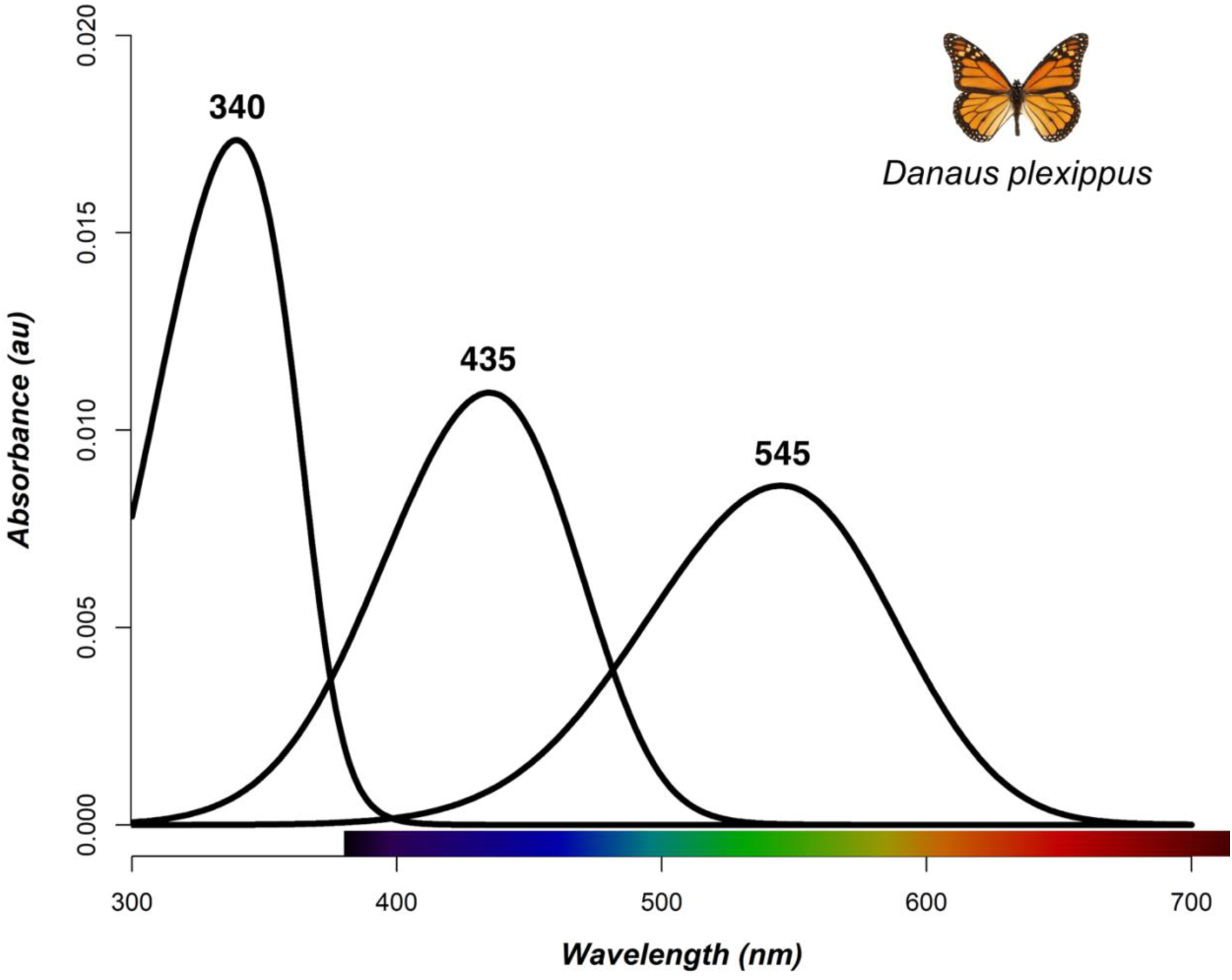
The butterfly visual model used to calibrate raw spectral irradiance values. Absorbance curves of the three visual pigments (from left to right: ultraviolet, blue, long-wavelength) in the adult compound eye of *Danaus plexippus* (Nymphalidae: Danainae), the most closely related species to ithomiines for which visual pigment sensitivities are known. The wavelengths of maximal absorbance (λmax) of these visual pigments are labelled above the peak of each curve and were used to calibrate our raw spectral irradiance readings for butterfly vision, using the template by Govardovskii *et al.* (*35*).

**Fig. S2.**
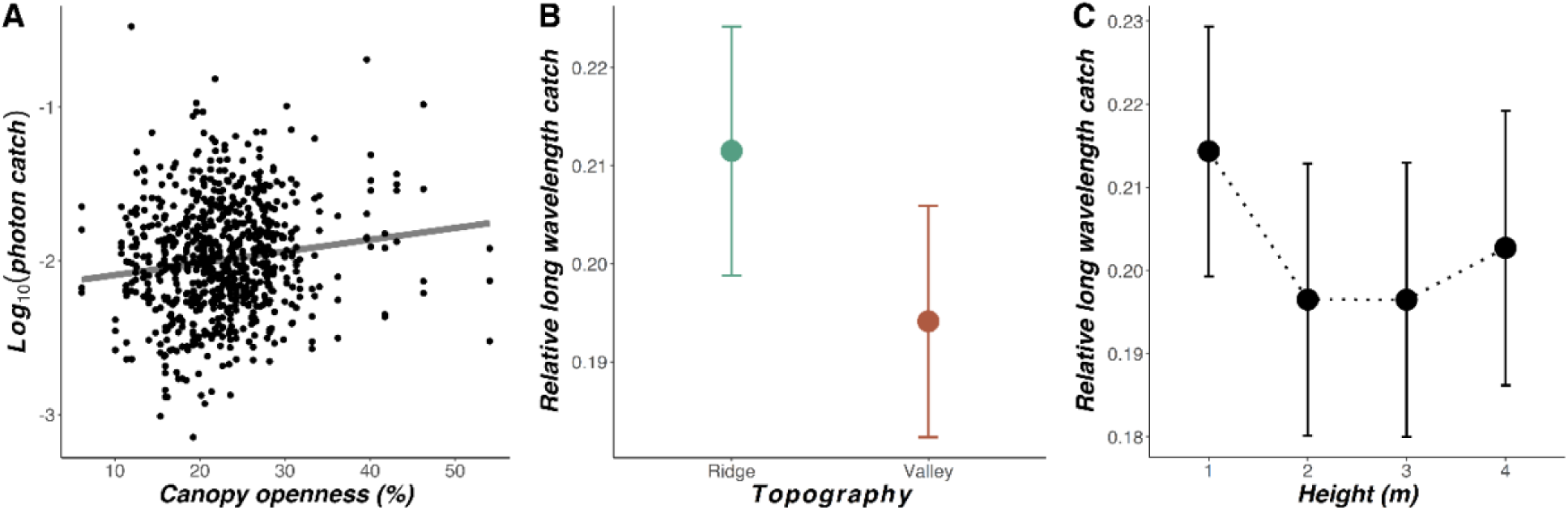
Additional ecological associations with light environment across a topographically variable forest transect (*N* = 762, 9 ridge/valley replicates). **(A,B)** Mean relative quantum catch of long wavelengths with respect to transect topography (A) and height from the ground (m) (B). Error bars represent 95% confidence intervals. **(C)** Relationship between (log10-transformed) photon catch (10^10^ quanta s^-1^ m^-2^) for the LW sensitivity function and canopy openness overlayed with the regression line estimated from linear mixed effect modelling.

**Fig. S3.**
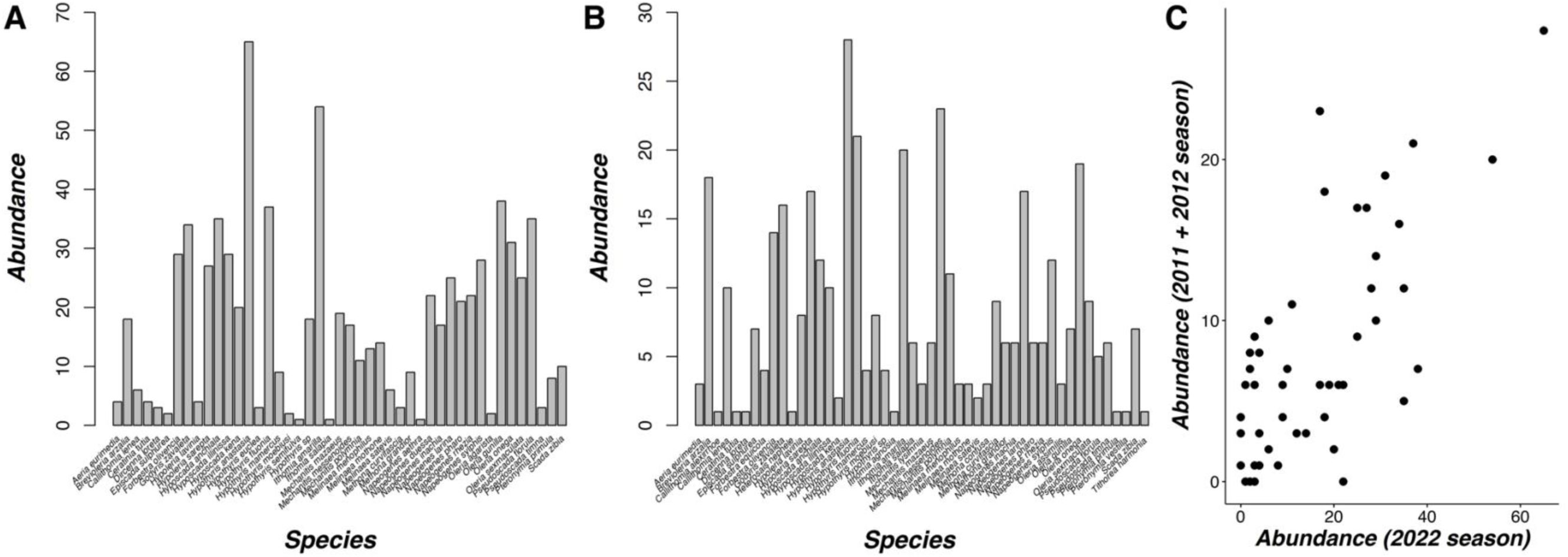
Ithomiini species abundances within the Ecuadorean community. **(A,B)** Histograms denoting the abundance distribution of species found in 2022 (*N* = 785, 45 species) (A), from which ecological and eye samples were obtained, and in 2011 and 2012 combined (*N* = 392, 49 species) (B), from which brain samples and wing morphological measurements were obtained. **(C)** The positive relationship between species abundance from the 2022 field season and the 2011/2012 field seasons combined.

**Fig. S4.**
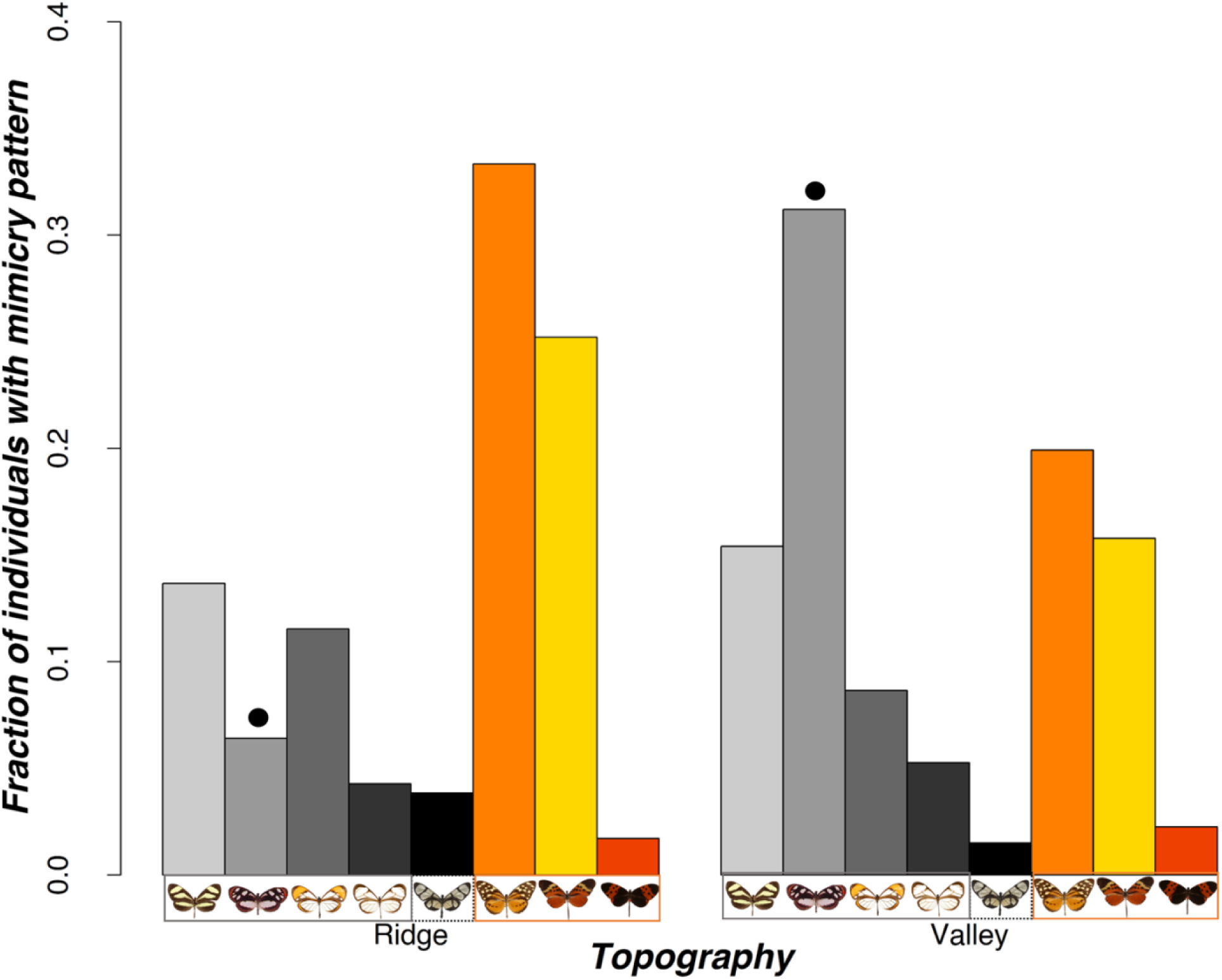
Relative mimicry ring abundance at ridge and valley sites along a topographically variable forest transect. Example models of each mimicry ring are shown below their corresponding bar, grouped based on their general color pattern classification. Each black dot represents a mimicry ring whose abundance significantly differed between ridges and valleys in the MCMCglmm analysis.

**Fig. S5.**
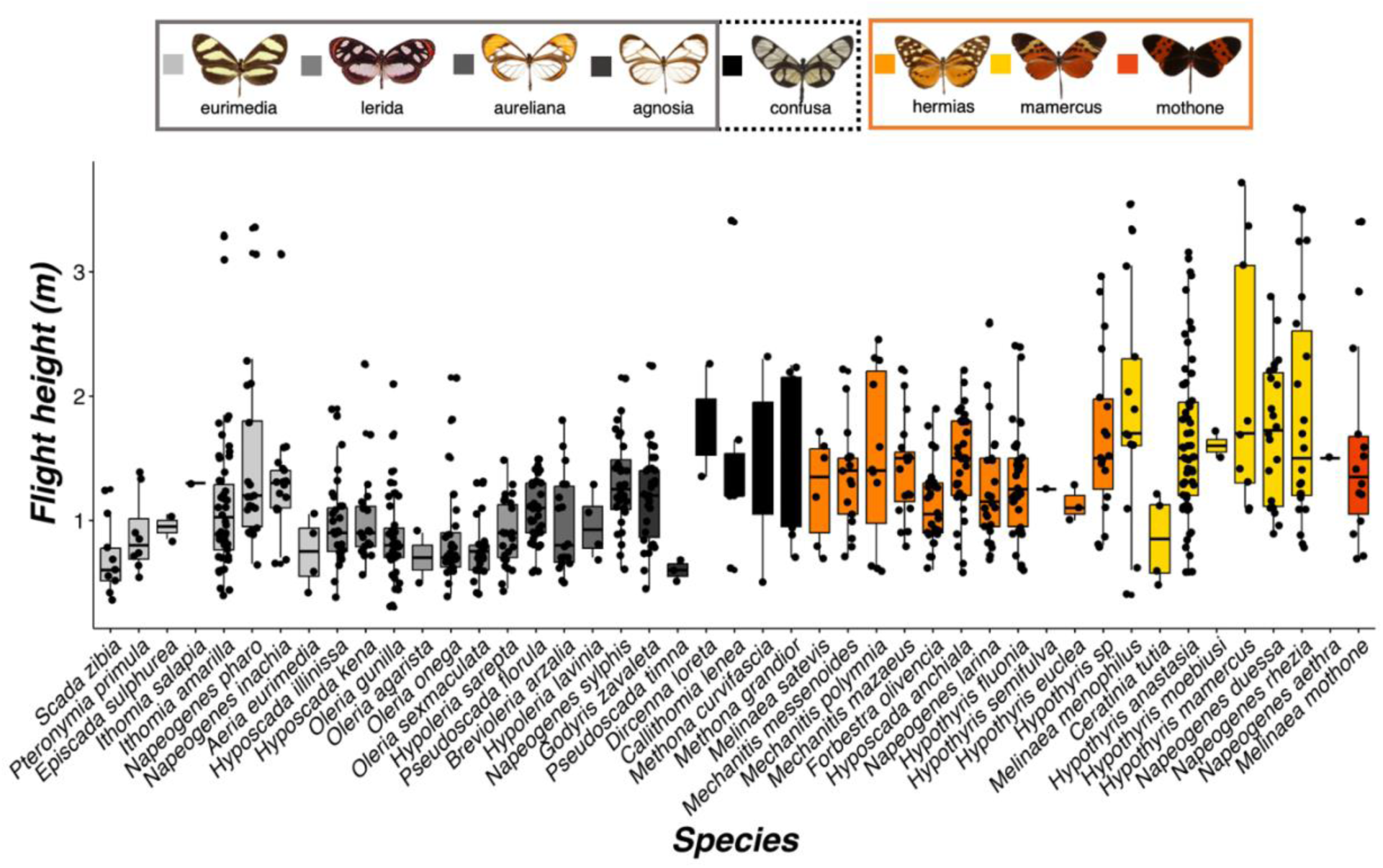
Flight height (m) differences between ithomiine species, coded by mimicry ring (*N* = 785, 45 species). Example models of each mimicry ring are shown on the top row, grouped based on their general color pattern classification. Medians (thick horizontal bars), interquartile ranges (boxes), values within 1.5 interquartile ranges of the box edges (whiskers), and possible outliers (datapoints outside whiskers) are plotted.

**Fig. S6.**
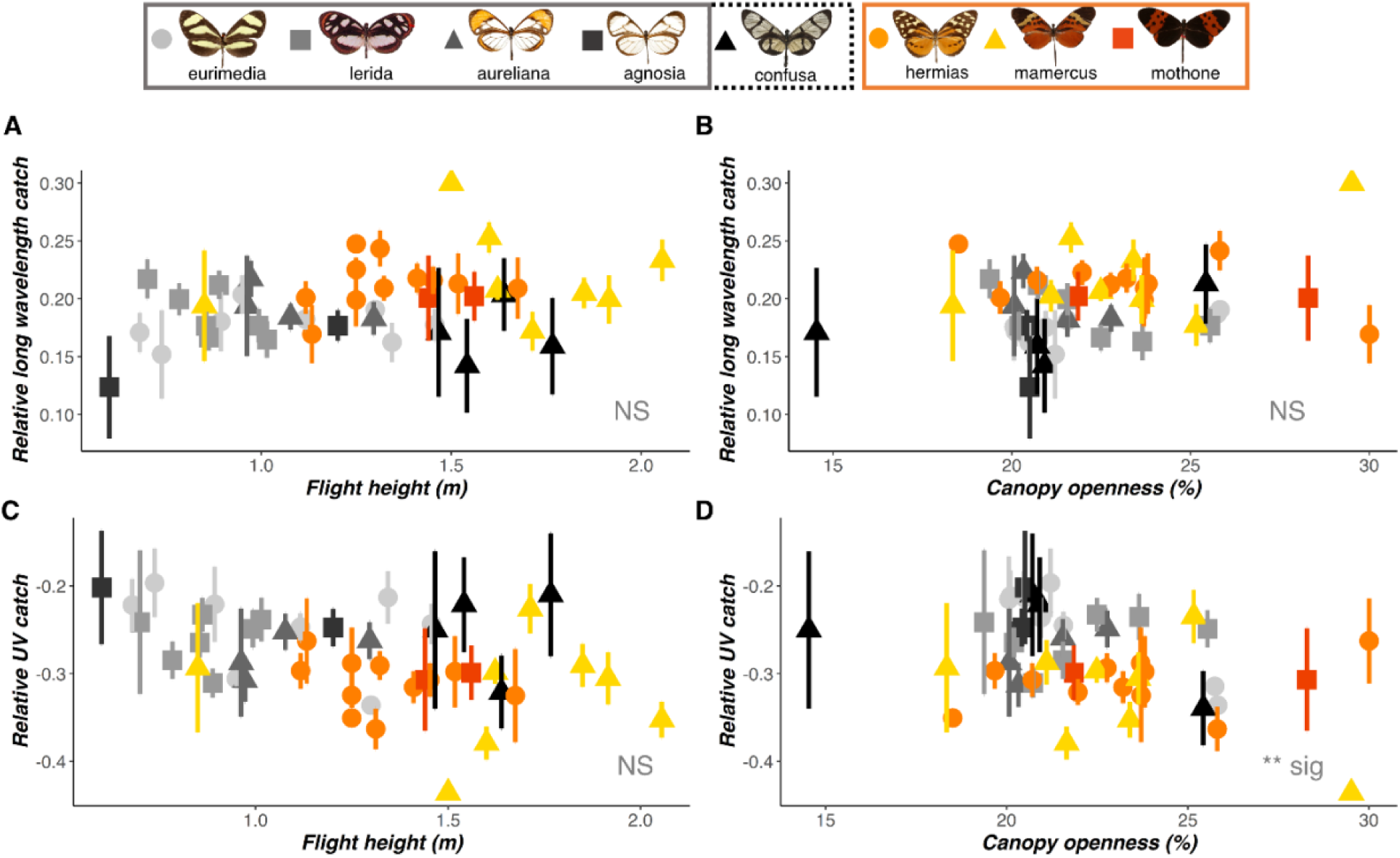
Variation in spectral composition between ithomiine microhabitats. **(A-D)** Mean relative quantum catch of long wavelengths (A,B) and ultraviolet (C,D) of each species (*N* = 785, 45 species), coded by mimicry ring, plotted against mean flight height (m; A,C) and canopy openness (%; B,D). Error bars indicate standard error. Mimicry rings were shown to significantly segregate with respect to the relative catch of long wavelengths, but not ultraviolet. Example models for each mimicry ring are shown on the top row, grouped based on their general color pattern classification. Significance of each ecological variable is indicated at the bottom right of each panel. NS P > 0.05, *P < 0.05, **P < 0.01, ***P < 0.001.

**Fig. S7.**
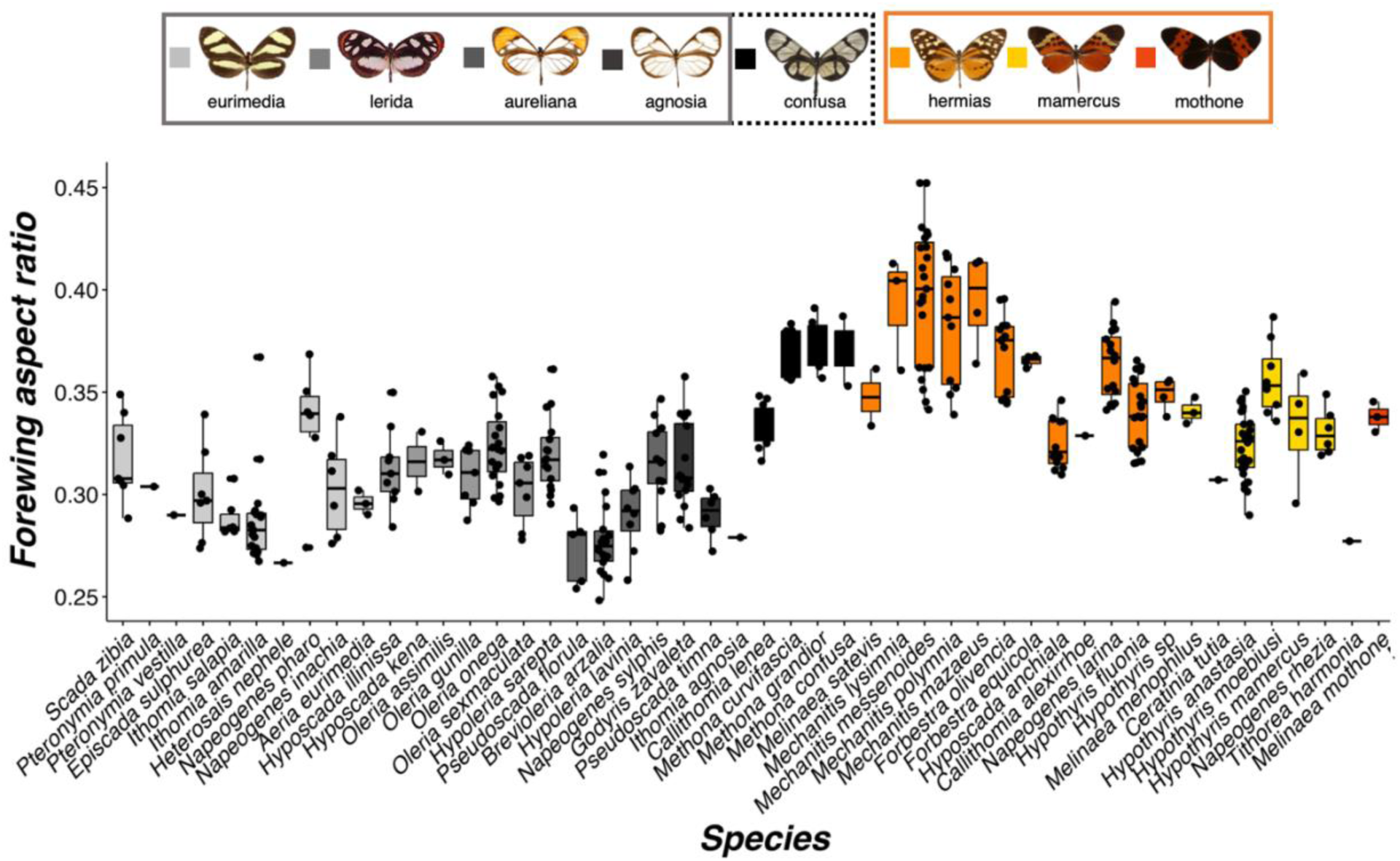
Forewing aspect ratio differences between ithomiine species, coded by mimicry ring (*N =* 392, 49 species). Example models of each mimicry ring are shown on the top row, grouped based on their general color pattern classification. Medians (thick horizontal bars), interquartile ranges (boxes), values within 1.5 interquartile ranges of the box edges (whiskers), and possible outliers (datapoints outside whiskers) are plotted.

**Fig. S8.**
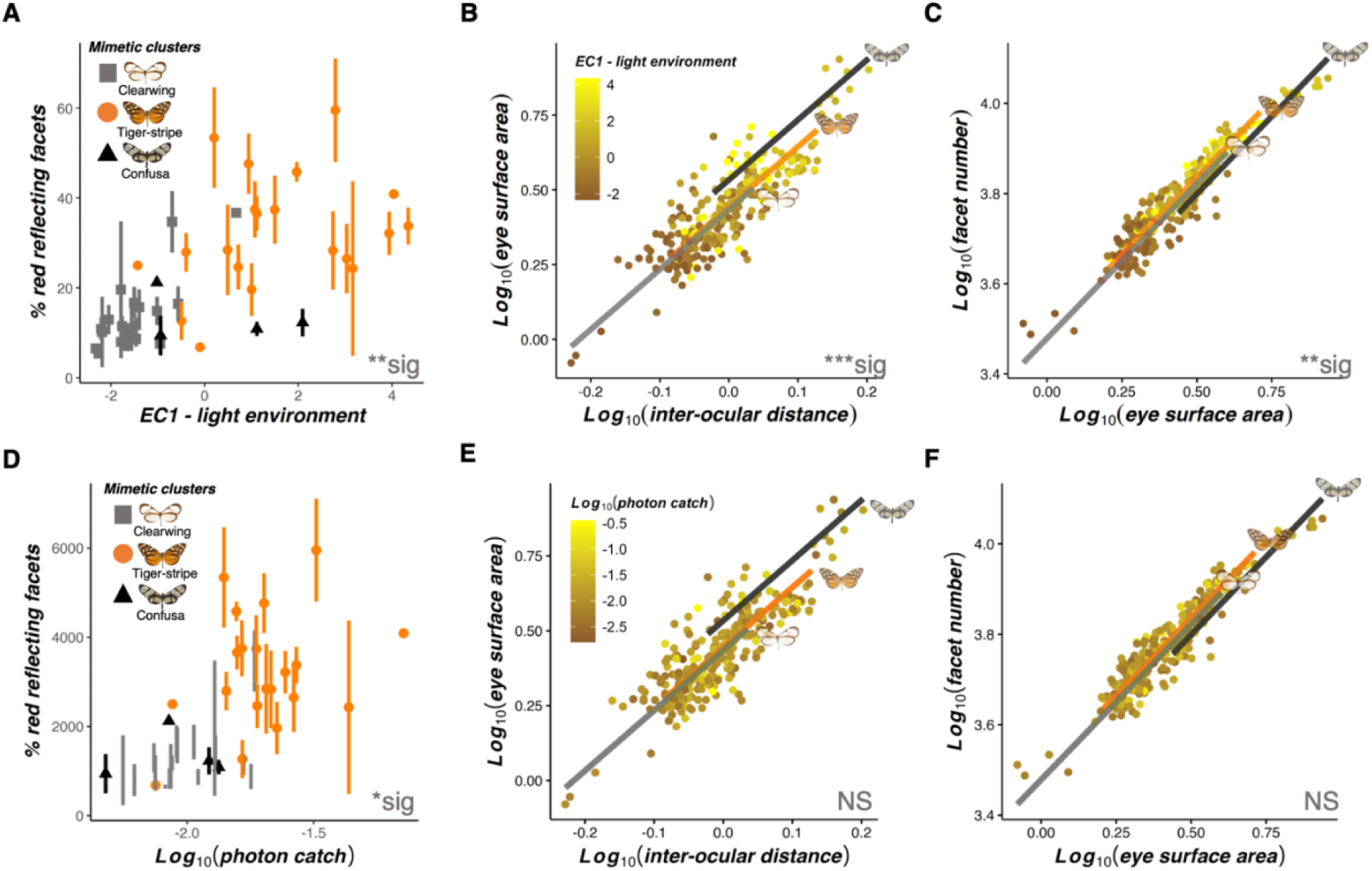
Additional eye physiological and anatomical associations with light environment and mimicry (*N =* 363, 45 species). **(A-F)** Correlation of traits with respect to EC1 for each species (A-C) and (log10-transformed) photon catch (10^10^ quanta s^-1^ m^-2^) for the LW sensitivity function (D-F), showing differences in the proportion of red reflecting facets (shown as species means coded by mimetic cluster) (A,D), eye surface area (mm^2^) when scaled against inter-ocular distance (mm) (B,E), and number of facets when scaled against eye surface area (C,F). For each panel containing an allometric control, all variables are log10-transformed and regression lines for each mimetic cluster, estimated from standardized major axis regression are superimposed on top, alongside example models (B,C,E,F). The color scale for EC1 and photon catch are shown on the top left of (B) and (E) respectively. Asterisks at the bottom right of each panel indicate the significance level of EC1 (A-C) and photon catch (D-F) at explaining variation in each trait. NS P > 0.05, *P < 0.05, **P < 0.01, ***P < 0.001.

**Fig. S9.**
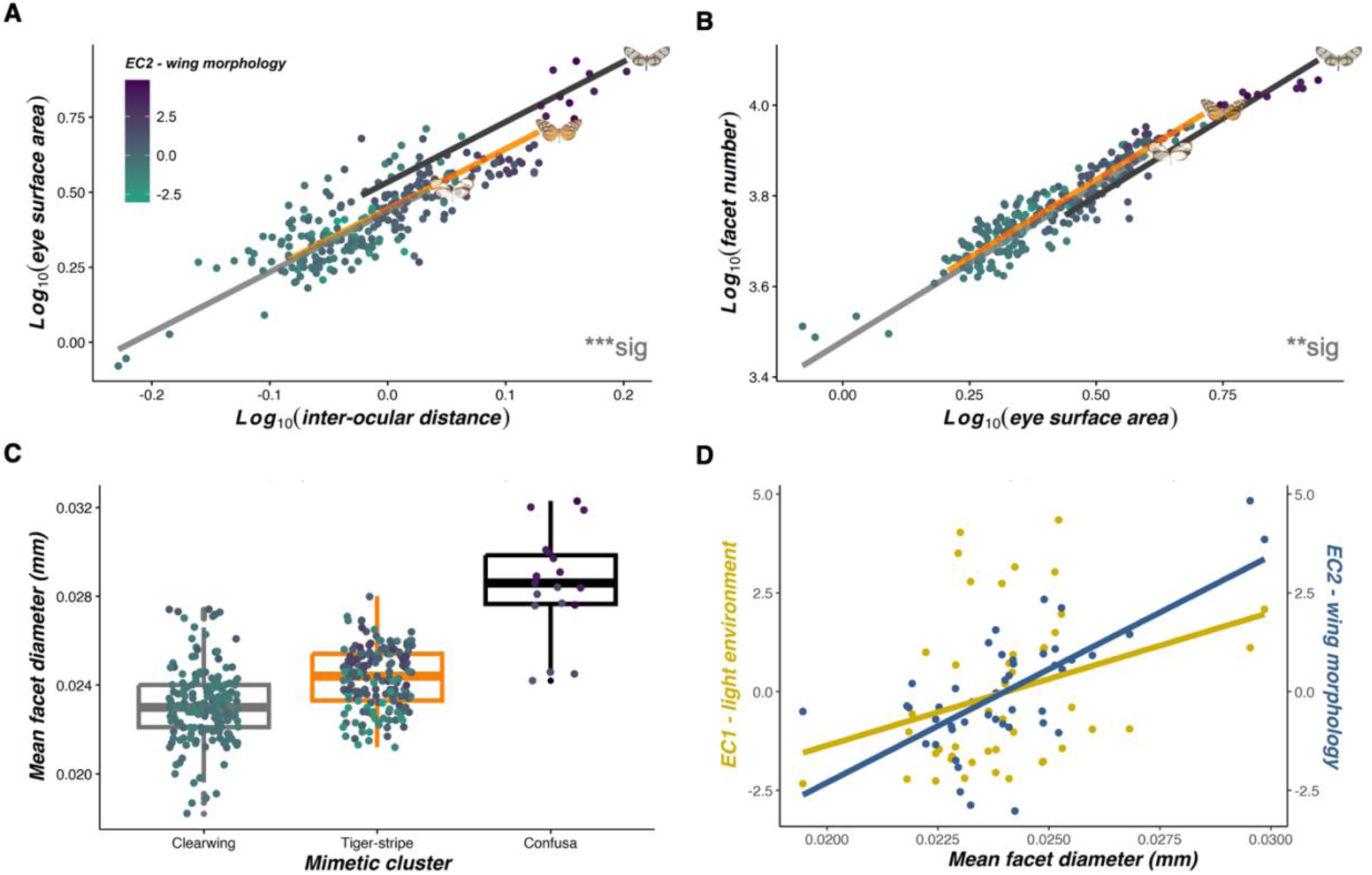
Eye anatomical associations with wing morphology (*N =* 363, 45 species). **(A,B)** Non-allometric convergence between EC2 for each species and eye surface area (mm^2^), when scaled against inter-ocular distance (mm) (A), and facet number, when scaled against eye surface area (B). All eye anatomy traits in (A,B) are log10-transformed. Regression lines for each mimetic cluster, estimated from standardized major axis regressions are superimposed on top, alongside example models. **(C)** Convergence in mean facet diameter (mm) for individuals with similar wing morphologies, separated by mimetic cluster. Medians (thick horizontal bars), interquartile ranges (boxes), values within 1.5 interquartile ranges of the box edges (whiskers), and possible outliers (datapoints outside whiskers) are plotted. In (A-C), turquoise-purple color shades represent the mean EC2 value for each species, the color scale for which is shown in the top left of (A). Asterisks at the bottom right of each panel indicate the significance level of EC2 at explaining variation in each trait. NS P > 0.05, *P < 0.05, **P < 0.01, ***P < 0.001. **(D)** Positive correlation between the mean facet diameter of each species and both ecological PC axes, representing light environmental (EC1, yellow) and wing morphological variation (EC2, blue).

**Fig. S10.**
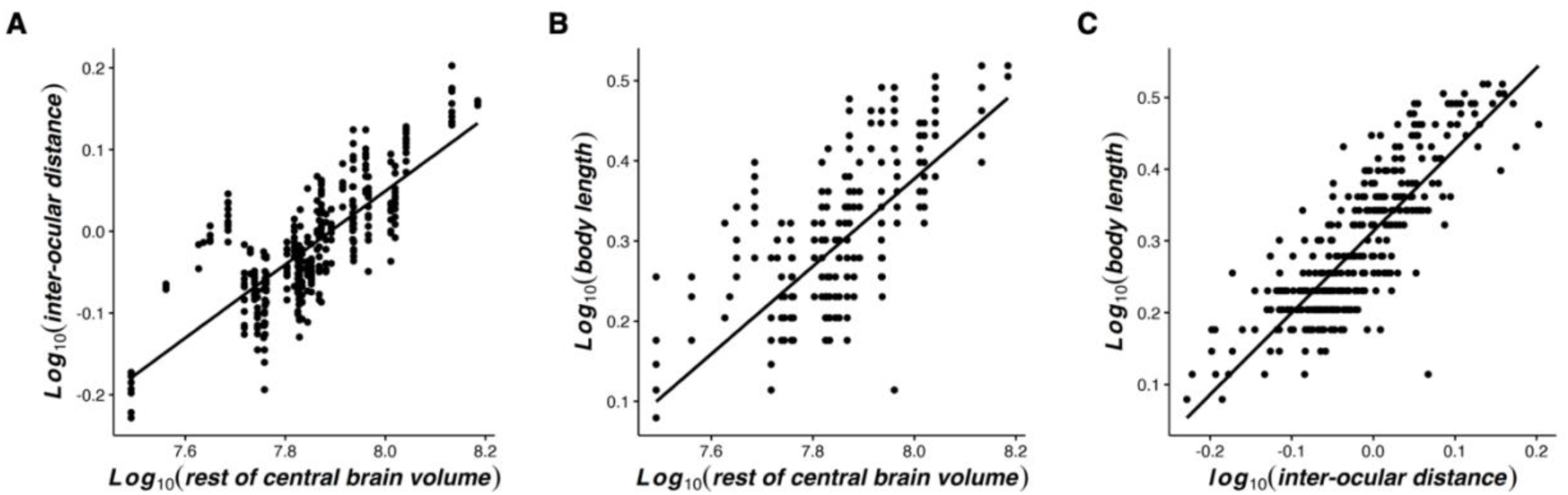
Positive correlations between log10-transformed allometric controls. **(A,B)** Inter-ocular distance (mm) (A) and body length (cm) (B) plotted against the mean “rest of central brain” volume (μm^3^) of each species. **(C)** Body length plotted against inter-ocular distance. All traits are log10-transformed.

**Fig. S11.**
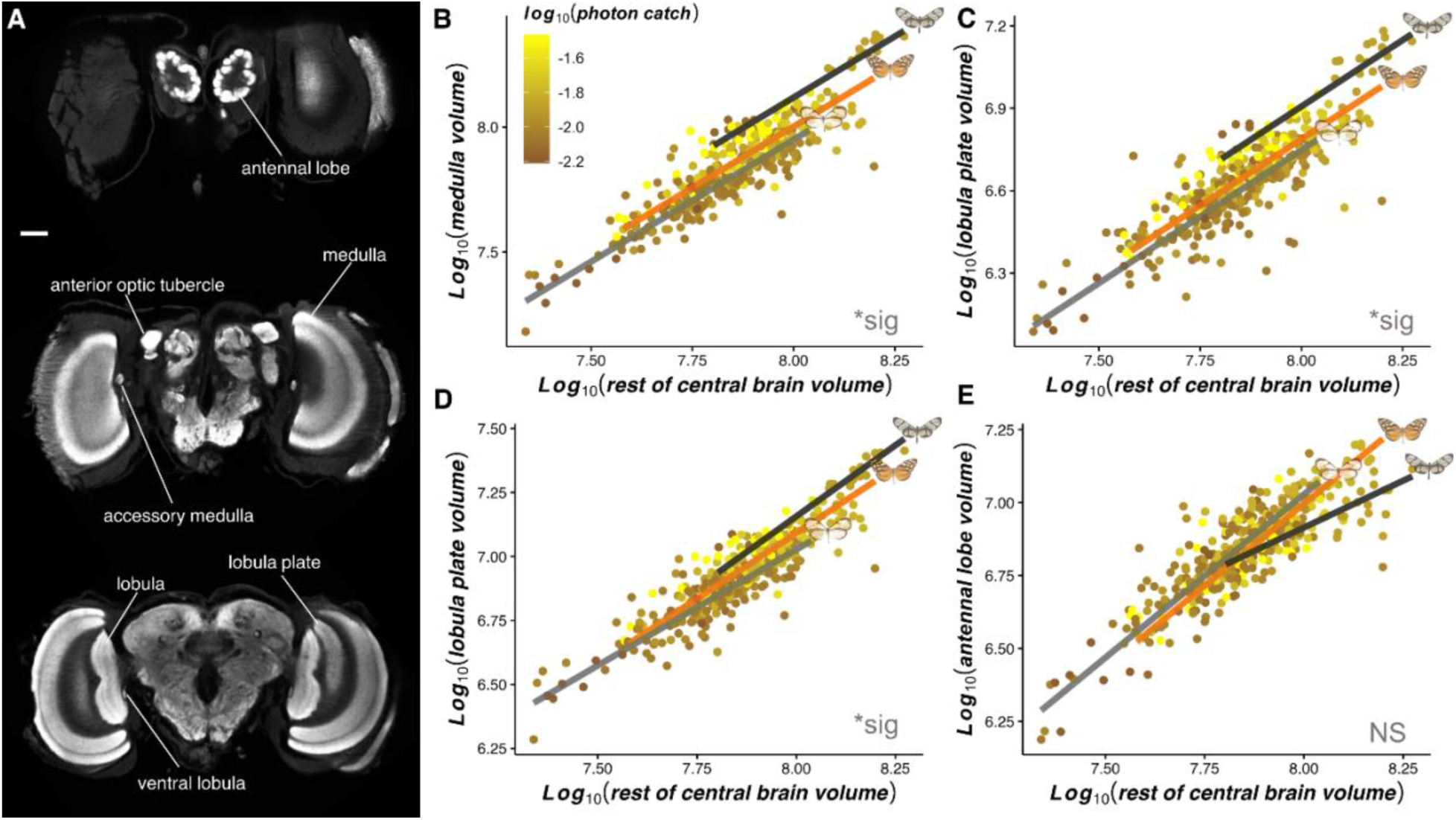
Additional neuroanatomical associations with light environment and mimicry (*N =* 374, 40 species). **(A)** Anti-synapsin immunofluorescence from frontal confocal brain sections of *Ithomia amarilla*, with imaging performed progressively along the posterior axis, moving from top to bottom. All reconstructed neuropils are labelled. Scale bar = 100 μm. **(B-E)** Non-allometric convergence between the mean overall photon catch (10^10^ quanta s^-1^ m^-2^) for the LW sensitivity function of each species and the level of volumetric investment (μm^3^) in the medulla (B), lobula plate (C), lobula (D), and antennal lobe (E) when scaled against the volume of the “rest of central brain”. Brown-yellow color shades represent the mean photon catch for each species, the color scale for which is shown in the top left of (B). All variables are log10-transformed. Regression lines for each mimetic cluster, estimated from standardized major axis regressions are superimposed on top, alongside example models. Asterisks at the bottom right of each panel indicate the significance level of photon catch at predicting relative investment in each neuropil. NS P > 0.05, *P < 0.05, **P < 0.01, ***P < 0.001.

**Fig. S12.**
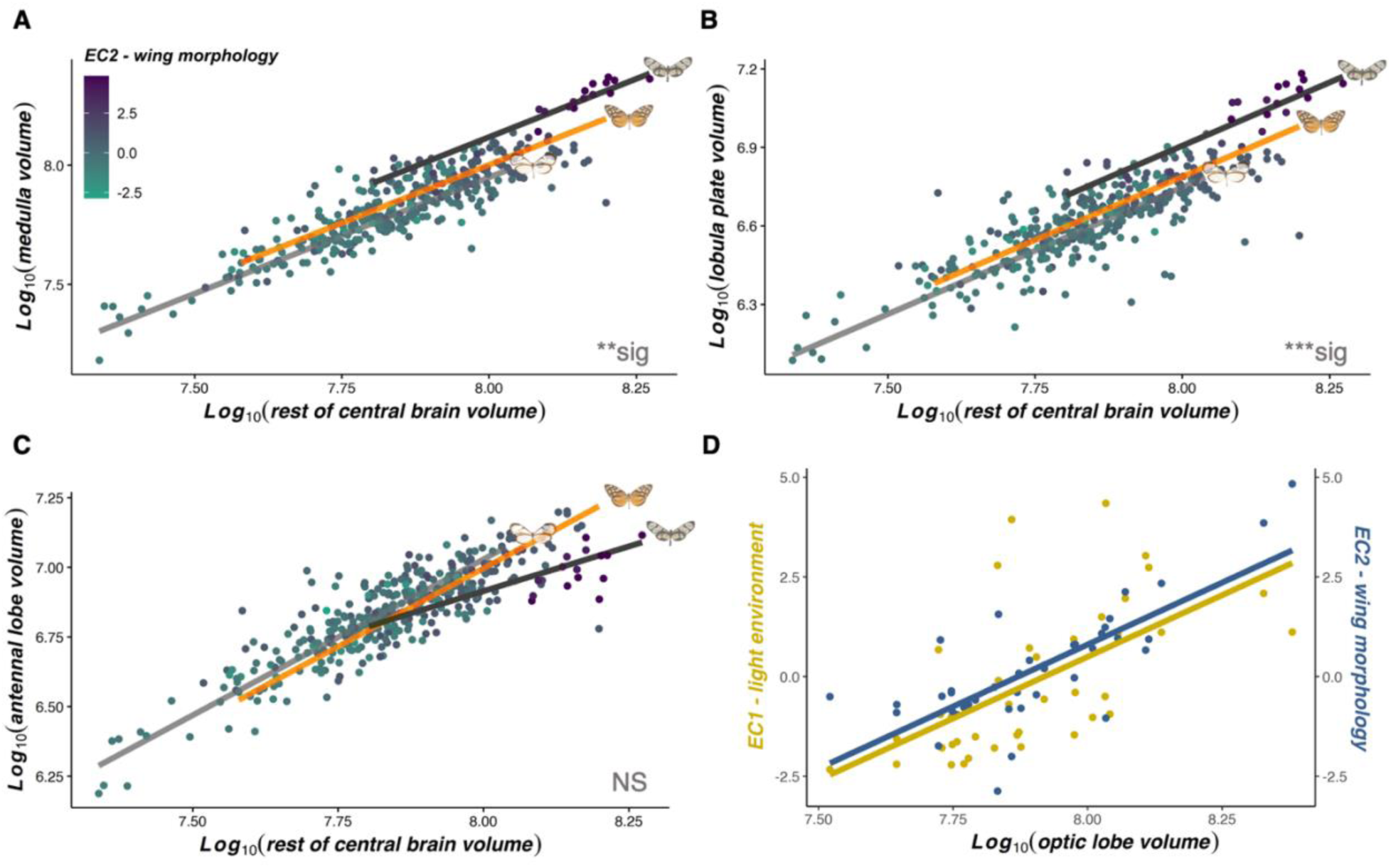
Neuroanatomical associations with wing morphology (*N =* 374, 40 species). **(A-C)** Non-allometric convergence between the mean wing morphology of each species and the volume (μm^3^) of the medulla (A), lobula plate (B), and antennal lobe (C), when scaled against the volume of the “rest of central brain”. All volumetric traits are log10-transformed. Turquoise-purple color shades represent the mean EC2 value for each species, the color scale for which is shown in the top left of (A). Regression lines for each mimetic cluster, estimated from standardized major axis regressions are superimposed on top, alongside example models. Asterisks at the bottom right of each panel indicate the significance level of EC2 at explaining variation in each neuropil. NS P > 0.05, *P < 0.05, **P < 0.01, ***P < 0.001. **(D)** Positive correlation between the mean gross optic lobe volume of each species and both ecological PC axes, representing light environmental (EC1, yellow) and wing morphological variation (EC2, blue).

**Fig. S13.**
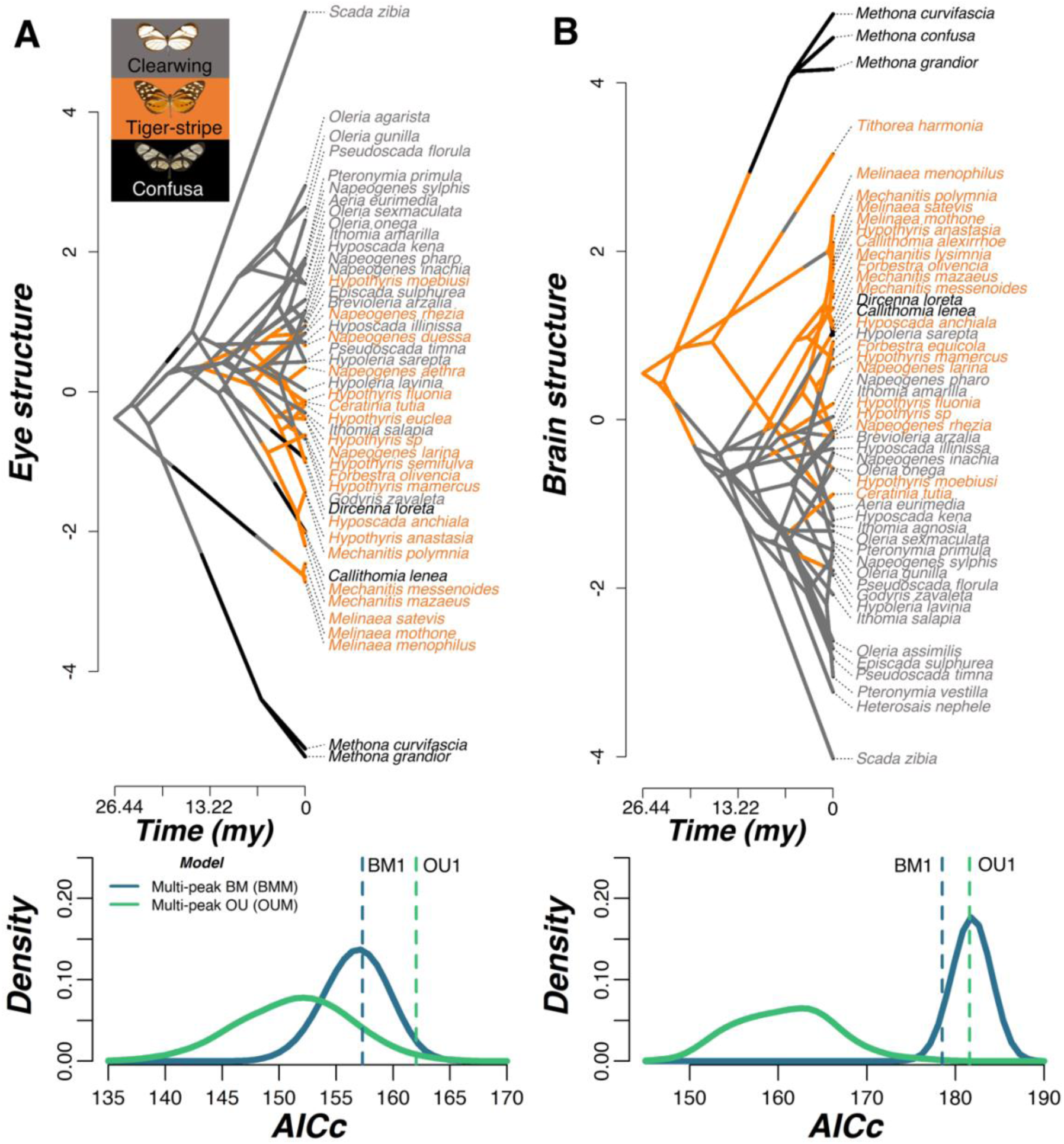
Convergence in eye and brain structure separately among co-mimics. **(A,B)** Above are phenograms based on PC1 of two principal component analyses which summarized variation in eye (*N* = 45 species) (A) and brain (*N* = 49 species) (B) structure based on traits which showed significant light environmental effects. Instances where branches cross and concentrate in a given area indicate convergent lineages. Below each phenogram are Kernel density plots of small sample corrected Akaike Information Criterion (AICc) scores obtained from multi-peak Brownian motion (BM) and Ornstein-Uhlenbeck (OU) evolutionary models, constructed from 500 simulated character maps where species belonging to the same mimetic cluster were assigned to the same selective regime. Blue and green vertical dashed lines indicate the AICc score for single-peak BM (BM1) and OU (OU1) models respectively. Example models of each mimetic cluster are shown in the top left of (A).

**Table S1 (separate file). Light microhabitat variation and partitioning along a topographically variable transect. (A)** Linear mixed model output showing the effect of canopy openness, height from the ground, and topography (and their interaction) at explaining variation in each spectral variable. **(B)** MCMCglmm output, including pairwise comparisons, showing how the abundance of ithomiine mimicry rings differs with respect to ecological variation along the transect. **(C)** Scores and loadings from a principal component analysis, included in the above analysis, conducted on estimated photon catches of each photoreceptor for spectral measurements taken along the transect

**Table S2A (separate file). Light microhabitat variation and partitioning for individually caught ithomiine butterflies – MCMCglmm and PGLS segregation analyses.** MCMCglmm (i) and PGLS (ii) output, including pairwise comparisons, showing the effect of canopy openness, flight height, and mimicry ring at explaining visual niche variation for each spectral variable.

**Table S2B (separate file). Light microhabitat variation and partitioning for individually caught ithomiine butterflies – spectral PCA and light-wing morphological associations.** (i) Scores and loadings from a principal component analysis, included in the above analysis, conducted on estimated photon catches of each photoreceptor for spectral measurements taken for individual butterflies. (ii) MCMCglmm and PGLS output showing the effect of wing morphological variables at explaining visual niche variation for each spectral variable.

**Table S3 (separate file). Ecological principal component analysis (PCA). (A)** MCMCglmm and **(B)** PGLS output, including pairwise comparisons, showing the effect of canopy openness, flight height, and mimicry ring at explaining variation in each wing morphological variable.

**Table S4A (separate file). Visual system associations with light environment and flight-related morphology – ecological PC axes.** MCMCglmm (i) and PGLS (ii) output showing the effect of EC1 (light environment) and EC2 (wing morphology) at explaining variation in each visual trait, alongside allometric controls where appropriate. (iii) MCMCglmm and PGLS output showing correlations between allometric controls.

**Table S4B (separate file). Visual system associations with light environment and flight-related morphology – spectral and wing morphological variables.** MCMCglmm (i,ii) and PGLS (iii, iv) output showing the effect of individual spectral variables (i,iii) and individual wing morphological variables (ii,iv) at explaining variation in each visual trait, alongside allometric controls where appropriate.

**Table S5 (separate file). Sensory convergence between co-mimics. (A)** MCMCglmm (i) and PGLS (ii) output, including pairwise comparisons, showing the effect of mimetic cluster at explaining variation in each visual trait that showed significant light environmental associations, alongside allometric controls where appropriate. **(B)** Output from standardized major axis regression analysis, including pairwise comparisons, testing for a non-allometric effect of mimetic cluster at explaining the scaling relationship of each visual trait, using individual (i) and species mean (ii) data.

**Table S6 (separate file). Visual system principal component analyses (PCA). (A-C)** Scores and loadings from PCAs summarizing variation in eye+brain structure (A), and eye (B) and brain (C) structure separately.

**Table S7 (separate file). Evolutionary modelling of convergence. (A)** Summarized output from single/multi-peak Brownian motion (BM), Ornstein-Uhlenbeck (OU) and early burst (EB) models, created in *mvMORPH*, for EC1, EC2, eye+brain structure, and eye and brain structure separately, with co-mimics assigned to different selective regimes. **(B)** Raw output from each *mvMORPH* model simulation (*N* = 500) **(C)** Convergence (C) indices from a *convevol* analysis, testing for convergence in eye+brain structure.

